# Priors and Payoffs in Confidence Judgments

**DOI:** 10.1101/703082

**Authors:** Shannon M. Locke, Elon Gaffin-Cahn, Nadia Hosseinizaveh, Pascal Mamassian, Michael S. Landy

## Abstract

Priors and payoffs are known to affect perceptual decision-making, but little is understood about how they influence confidence judgments. For optimal perceptual decision-making, both priors and payoffs should be considered when selecting a response. However, for confidence to reflect the probability of being correct in a perceptual decision, priors should affect confidence but payoffs should not. To experimentally test whether human observers follow this normative behavior, we conducted an orientation-discrimination task with varied priors and payoffs, probing both perceptual and metacognitive decision-making. We then examined the placement of discrimination and confidence criteria according to several plausible Signal Detection Theory models. In the normative model, observers use the optimal discrimination criterion (i.e., the criterion that maximizes expected gain) and confidence criteria that shift with the discrimination criterion that maximizes accuracy (i.e., are not affected by payoffs). No observer was consistent with this model, with the majority exhibiting non-normative confidence behavior. One subset of observers ignored both priors and payoffs for confidence, always fixing the confidence criteria around the neutral discrimination criterion. The other group of observers incorrectly incorporated payoffs into their confidence by always shifting their confidence criteria with the same gains-maximizing criterion used for discrimination. Such metacognitive mistakes could have negative consequences outside the laboratory setting, particularly when priors or payoffs are not matched for all the possible decision alternatives.

## 2 Introduction

In making a perceptual decision, it is wise to consider information beyond the available sensory evidence. To maximize expected gains, one should consider both the baseline probability of each possible world state, i.e., *priors*, as well as the associated risks and rewards for choosing or not choosing each response alternative, i.e., *payoffs*. In the Signal Detection Theory (SDT) framework, priors and payoffs alter the threshold amount of evidence required to choose one alternative versus another, that is, a shift in the *criterion* for reporting option “A” versus option “B” in a binary task. For example, consider a radiologist trying to detect a tumor in an x-ray image. The radiologist should be more likely to report a positive result for a suspicious shadow if the patient’s file indicates they are a smoker, as this means they have a higher prior probability of cancer. Similarly, the high cost of waiting to treat the cancer should also bias the radiologist towards declaring a positive result. In both real and laboratory environments, observers have been found to factor in priors and payoffs to varying extents when setting the decision criterion (Maddox and Bohil, 1998, 2000; Maddox and Dodd, 2001; Wolfe et al., 2005; Ackermann and Landy, 2015; Horowitz, 2017).

Decisions about the state of the world (cancer or not cancer, cat or dog, tilted clockwise or counter-clockwise) are referred to as stimulus-conditioned responses or *Type 1* decisions. Judgments can also be made about our Type 1 decisions, such as our confidence in the decision, which are referred to as response-conditioned responses or *Type 2* decisions (Clarke et al., 1959; Galvin et al., 2003; Mamassian, 2016). Confidence judgments are often operationalized in binary decision-making experiments as a subjective estimate of the probability the Type 1 response was correct (Pouget et al., 2016). Confidence plays a broad role in guiding behavior, subsequent decision-making, and learning in a multitude of scenarios for both humans and animals (**?**Smith et al., 2003; Beran et al., **2012)**.

How does an ideal-observer radiologist modify confidence judgments in response to varying priors or payoffs? Intuitively, a radiologist should be more confident in a positive diagnosis when the patient is a smoker, given that they have been educated on the prior scientific evidence on the health risks of smoking. Additional confirmatory information should boost confidence in that positive diagnosis, and contrary evidence should reduce confidence, because priors (smoker or non-smoker) and sensory evidence (cancerous-looking shadow) are both informative about the likelihood over possible world states. However, this is not the case for payoffs. Incentivizing the different responses with rewards or punishment (e.g., delivering good or bad news) does not change the uncertainty about the world state. The radiologist should not be more or less confident in their cancer diagnosis if the type of cancer would be deadly or benign or if the surgical procedure is expensive or not, even though these factors should affect their initial diagnosis. In fact, sometimes payoffs will lead the decision-maker to choose the less probable alternative and this should be reflected by low confidence in the decision, such as the radiologist erring on the side of caution for a patient who smokes with an otherwise normal shadow on the x-ray image.

The literature is scarce on the issue of whether and how human observers adjust confidence in response to prior-payoff structures. In one perceptual study, the prior probabilities of target present versus absent affected the placement of the criteria for Type 1 and 2 judgments (Sherman et al., 2015), with some evidence that confidence better predicts performance for responses congruent with the more probable outcome than those that are incongruent. In the realm of social judgments, prior probabilities have been shown to modulate the degree of confidence, with higher confidence assigned to more probable outcomes (Manis et al., 1980). However, others have found counter-productive incorporation of priors, with over-confidence for low-probability outcomes and under-confidence for high-probability outcomes (Dunning et al., 1990). In regards to payoffs, early work on monetary incentives in perceptual categorization did collect confidence ratings, however they were not included in any analyses (Lee and Zentall, 1966). Consideration of payoff structures is ubiquitous in animal studies of confidence that employ post-decisional wagering methods (Smith et al., 2003). For example, in the opt-out paradigm, the animal is offered a choice between a small but certain reward and a risky alternative with either high reward or no reward, for correct and incorrect perceptual responses respectively. The fact that the animal chooses the small but certain reward in difficult trials is taken as evidence that it can distinguish between low and high levels of confidence (Kiani and Shadlen, 2009). However, because animals are motivated by expected gain and not explicit verbal instructions, it is impossible to isolate decision confidence that has not been confounded with the subjective value of the reward. That is, placing a monetary incentive on the confidence decision may alter the perceptual response.

We sought to characterize how human observers adjust their perceptual decisions and confidence in response to joint manipulation of priors and payoffs within the same perceptual task. We placed our participant in a visual orientation-discrimination task by presenting oriented Gabor patterns tilted left or right of vertical with a fixed orientation magnitude. In separate sessions we adjusted the prior-payoff structure by selecting the probability of a leftward-tilted versus rightward-tilted Gabor and by assigning different rewards for each of the response alternatives. We considered three classes of confidence behavior in our modeling. In the *normative-shift* models, priors but not payoffs determine the placement of confidence criteria; as discussed above, this is what participants should theoretically aim for. In the *gains-shift* models, both priors and payoffs determine confidence criteria; this is what would happen if participants ignored the reason why they shifted the first-order criterion. Finally, in the *neutral-fixed* models, the observer is insensitive to the prior-payoff context when placing confidence criteria. We also considered the possibility that participants were not optimal in using priors and payoffs in the discrimination decision. Therefore, variants of models within each class included 1) the nature of Type 1 criterion placement relative to optimal (e.g., decision conservatism), and 2) whether Type 1 conservatism was also present when participants were making their Type 2 decision. We found that almost all observers were best fit by either a gains-shift model or neutral-fixed model, neither of which constituted normative confidence behaviour. Furthermore, all observers who shifted the confidence criteria in response to changes in priors/payoffs maintained their Type 1 conservatism at the Type 2 metacognitive stage of decision-making. These results demonstrate a profound inability of our observers to correctly handle both priors and payoffs for metacognitive decision-making.

## 3 The Decision Models

Before presenting the outcome of our experiment, we describe the rationale and background for the modeling of Type 1 and Type 2 decision-making. This will allow us to directly interpret our behavioral results. We follow the example of a left-right orientation judgment followed by a binary low-high confidence judgment to match the experimental paradigm used in the present study. First the range of Type 1 models are identified, which assess the placement of the discrimination-decision criterion under different prior-payoffs scenarios. Then the Type 2 models are outlined, describing the different potential relationships between the decision criteria for confidence and the criterion for discrimination.

### 3.1 The Type 1 Decision

To make the Type 1 decision, observers must relate a noisy internal measurement, *x*, of the stimulus, *s*, where *s* ∈ {*s*_*L*_, *s*_*R*_}, to a binary response, which in the context of our experiment is “tilted left” (say “*s* = *s*_*L*_”) or “tilted right” (say “*s* = *s*_*R*_”). This is done by a comparison to an internal criterion, *k*_1_, such that if *x* < *k*_1_, the observer will respond with”tilted left”, and otherwise “tilted right” (Figure 1a). The only component of the Type 1 model the observer controls is the placement of the criterion. The optimal value of *k*_1_ (*k*_*opt*_) maximizes the expected gain, ensuring the observer makes the most points/money/etc. over the course of the experiment. The value of *k*_*opt*_ depends on three things:

**Figure 1:**
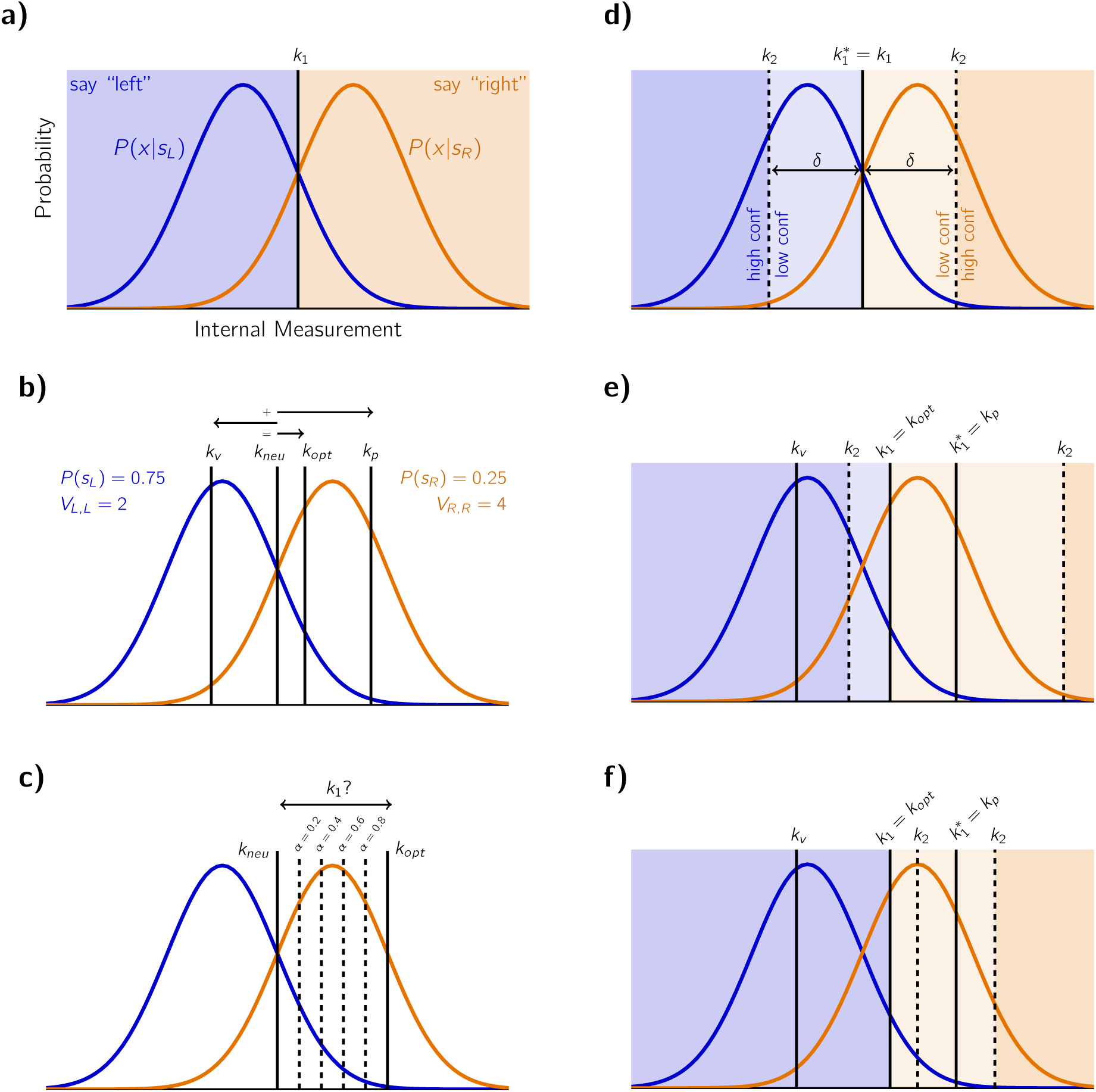
Illustration of the full SDT model. a) On each trial, an internal measurement of stimulus orientation is drawn from a Gaussian probability distribution conditional on the true stimulus value. The Type 1 criterion, *k*_1_, defines a cut-off for reporting “left” or “right”. The ideal observer in a symmetrical priors and payoffs scenario is shown. b) The ideal observer’s criterion placement with both prior and payoff asymmetry. This prior asymmetry encourages a rightward criterion shift to *k*_*p*_ and the payoff asymmetry a leftward shift to *k*_*v*_. The optimal criterion placement that maximizes expected gain, *k*_*opt*_, is a sum of these two criterion shifts. For comparison, the neutral criterion, *k*_*neu*_ is shown. As the prior asymmetry is greater than the payoff asymmetry, 3:1 vs 1:2, *k*_*opt*_ ≠ *k*_*neu*_. c) A sub-optimal conservative observer will not adjust their Type 1 criterion far enough from *k*_*neu*_ to be optimal. The parameter *α* describes the degree of conservatism, with values closer to 0 being more conservative and closer to 1 less conservative. d) In the case of symmetric payoffs and priors, the Type 2 confidence criteria, *k*_2_, are placed equidistant from the Type 1 decision boundary by ±*δ*, carving up the internal measurement space into a low- and high-confidence region for each discrimination response option. e) For the normative Type 2 model, the confidence criteria are placed symmetrically around a hypothetical Type 1 criterion that only maximizes accuracy 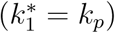. This figure shows the division of the measurement space as per the prior-payoff scenario in (b). As a left-tilted stimulus is much more likely, this results in many high-confidence left-tilt judgments and few high-confidence right-tilt judgments. Note that left versus right judgments still depend on *k*_1_. f) The same as in (e) but with a small value of *δ*. Note the low-confidence region where confidence should be high (left of the left-hand *k*_2_). This happens because in this region the observer will choose the Type 1 response that conflicts with the accuracy-maximizing criterion, hence they will report low confidence in their decision. Note that the displacements of the criteria from the neutral criterion in this figure are exaggerated for illustrative purposes.

i. The sensitivity of the observer, *d*′. In the standard model of the decision space, *P* (*x*|*s*_*L*_) ∼ *N* (*µ*_*L*_, *σ*_*L*_) and *P* (*x*|*s*_*R*_) ∼ *N* (*µ*_*R*_, *σ*_*R*_), with *µ*_*L*_ = −*µ*_*R*_ and *σ*_*L*_ = *σ*_*R*_ = 1. Under this transformation, the sensitivity *d*′ corresponds to the distance between the peaks of the two internal measurement distributions.
ii. The prior probability of each stimulus alternative, *P* (*s*_*L*_) and *P* (*s*_*R*_) = 1 − *P* (*s*_*L*_).
iii. The rewards for the four possible stimulus-response pairs, *V*_*r,s*_, which are the rewards (positive) or costs (negative) of responding *r* when the stimulus is *s*.

An ideal observer that maximizes expected gain (Green and Swets, 1966) uses criterion

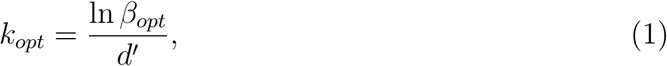

where the likelihood ratio *β*_*opt*_ at the optimal criterion is a function of priors and payoffs:

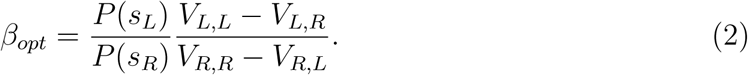

In our experiment, 0 points are awarded for incorrect answers, allowing us to simplify:

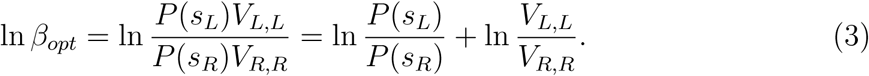

Thus, *k*_*opt*_ = *k*_*p*_ + *k*_*v*_, where *k*_*p*_ is the optimal criterion location if only priors were asymmetric and *k*_*v*_ is the optimal criterion if only the payoffs were asymmetric. As can be seen in Eq. 3, the effects of priors and payoffs sum when determining the optimal criterion (illustrated in Figure 1b). When the priors are more similar, or the payoffs are closer to equal, *k*_*opt*_ is closer to the neutral criterion *k*_*neu*_ = 0. Note that in the case of symmetric payoffs, *k*_*opt*_ maximizes both expected gain and expected accuracy, whereas when asymmetric payoffs are involved, *k*_*opt*_ maximizes expected gain only (i.e., *k*_*opt*_ ≠ *k*_*p*_). This is because to maximize expected gain, from time to time the observer is incentivized to choose the less probable outcome because it is more rewarded.

### 3.2 Conservatism

Often, human observers use a sub-optimal value of *k*_1_ when the prior probabilities or payoffs are not identical for each alternative. A common observation is that the criterion is not adjusted far enough from the neutral criterion towards the optimal criterion, *k*_*neu*_ < *k*_1_ < *k*_*opt*_ or *k*_*neu*_ > *k*_1_ > *k*_*opt*_, a behavior referred to as conservatism (Green and Swets, 1966; Maddox, 2002). It is useful to express conservatism as a weighted sum of the neutral and optimal criterion:

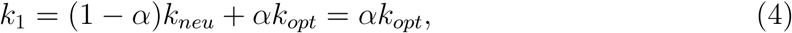

with 0 < *α* < 1 indicating conservative criterion placement. The degree of conservatism is greater the closer *α* is to 0 (Figure 1c). Several studies have contrasted the conservatism for unequal priors versus unequal payoffs, typically finding greater conservatism for unequal payoffs (Lee and Zentall, 1966; Ulehla, 1966; Healy and Kubovy, 1981; Ackermann and Landy, 2015) with few exceptions (Healy and Kubovy, 1978). This may result from an underlying criterion-adjustment strategy that depends on the shape of the expected-gain curve (as a function of criterion placement) and not just on the position of the optimal criterion maximizing expected gain (Busemeyer and Myung, 1992; Ackermann and Landy, 2015) or a strategy that trades off between maximizing expected gain and maximizing expected accuracy (Maddox, 2002; Maddox and Bohil, 2003). Given that the effects of priors and payoffs sum in Eq. 3, we will consider a sub-optimal model of criterion placement that has separate conservatism factors for payoffs and priors:

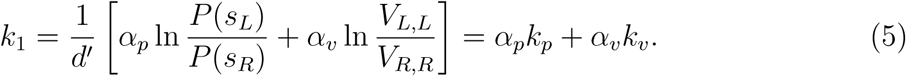

The conservatism factors, *α*_*p*_ and *α*_*v*_, scale these individually before they are summed to give the final conservative criterion placement, taking into account both prior and payoff asymmetries. This formulation allows for differing degrees of conservatism for priors and payoffs.

### 3.3 Type 1 Decision Models

We consider four models of the Type 1 discrimination decision in this paper, including the optimal model (i) and three sub-optimal models that include varying forms of conservatism (ii-iv):

i. Ω_1,*opt*_ : *k*_1_ = *k*_*opt*_ = *k*_*p*_ + *k*_*v*_
ii. Ω_1,1*α*_ : *k*_1_ = *αk*_*opt*_ = *α* (*k*_*p*_ + *k*_*v*_)
iii. Ω_1,2*α*_ : *k*_1_ = *α*_*p*_*k*_*p*_ + *α*_*v*_*k*_*v*_
iv. 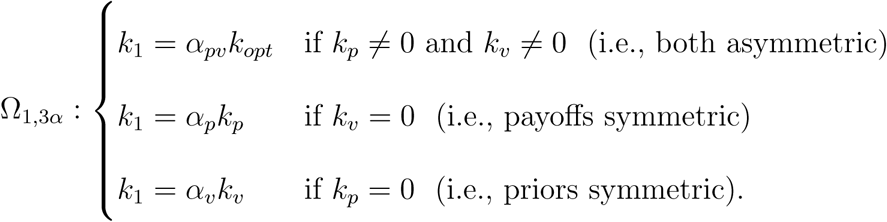

Thus, we consider models with no conservatism (Ω_1,*opt*_), with an identical degree of conservatism due to asymmetric priors and payoffs (Ω_1,1*α*_), or different amounts of conservatism for prior versus payoff manipulations (Ω_1,2*α*_). In the fourth model, we drop the assumption (that was based on the optimal model) that effects of payoffs and priors on criterion sum, i.e., that behavior with asymmetric priors and payoffs can be predicted from behavior with each effect alone (Ω_1,3*α*_). We consider this final model because the additivity of criterion shifts (Eq. 3) has not yet been experimentally confirmed with human observers (Stevenson et al., 1990).

In all models, we also consider an additive bias term, *γ*, corresponding to a perceptual bias in perceived vertical. The bias is also included in the neutral criterion *k*_*neu*_ = *γ*. For clarity, however, we have omitted it from the mathematical descriptions of the models. Note that any observer best fit by Ω_1,*opt*_ but with a *γ* significantly different from 0 would no longer be considered as having optimal behavior.

### 3.4 Confidence Criteria

Confidence judgments should reflect the belief that the selected alternative in the discrimination decision correctly matches the true world state. Generally speaking, the further the internal measurement is from a well-placed decision boundary, the more evidence there is for the discrimination judgment. This is instantiated in the extended SDT framework by the addition of two or more confidence criteria, *k*_2_ (Maniscalco and Lau, 2012, 2014). There are two such criteria for a binary confidence task and more confidence criteria when more than two confidence levels are provided. We restrict our treatment to the binary case, which can be trivially extended to include more gradations of confidence.

As illustrated in Figure 1d, for the case of symmetric payoffs and priors, there is a *k*_2_ confidence criterion on each side of the *k*_1_ decision boundary. If the measurement obtained is beyond one of these criteria relative to *k*_1_, then the observer will report high confidence, and otherwise will report low confidence. Stated another way, the addition of the confidence criteria effectively divides the measurement axis into four regions: high-confidence left, low-confidence left, low-confidence right, and high-confidence right. The closer to the discrimination decision boundary that the observer places *k*_2_, the more high-confidence responses they will give. We denote this distance as *δ. δ* is not always assumed to be identical for both confidence criteria (e.g. Maniscalco and Lau, 2012), but we assumed a single value of *δ* for model simplicity. Type 2 judgments were not incentivized in our experiment to allow observers to make a discrimination decision that was not influenced by a monetary reward on the confidence decision. Thus, there is no explicit cost function to constrain the distance parameter *δ*, so the precise setting of *δ* will not factor into the evaluation of how well the normative model fits observer behavior.

### 3.5 The Counterfactual Type 1 Criterion

The above description of how confidence responses are generated is well suited to cases where the payoffs are symmetric. This is because the optimal Type 1 decision criterion maximizes both gain and accuracy. For an internal measurement at the discrimination boundary, it is equally probable that the stimulus had a rightward versus leftward orientation. Expressed another way, the log-posterior ratio at *k*_*opt*_ is 1. Thus, the distance from the discrimination boundary is a good measure for the probability that the Type 1 response is correct (i.e., confidence as we defined it above). This, however, is not the case when payoffs are asymmetric (*k*_1_ = *k*_*p*_ + *k*_*v*_ = *k*_*opt*_ where *k*_*v*_ ≠ 0), as the ideal observer maximizes gain but not accuracy. The log-posterior ratio is not 1 at *k*_*opt*_ but rather it is equal to 1 at *k*_*p*_.

To extend the SDT model of confidence to asymmetric payoffs, we introduce a new criterion. We call counterfactual criterion, 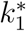, the criterion that the ideal observer would have used if they ignored the payoff structure of the environment and exclusively maximized accuracy and not gain (i.e., 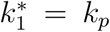). It is this discrimination criterion that confidence criteria are yoked to in our normative model (Figure 1e). Note that whenever payoffs are symmetrical (*k*_*v*_ = 0), 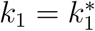. Figure 1f illustrates a situation unique to this model that may occur when payoffs are asymmetric. Here, the value of *δ* is sufficiently small that both *k*_2_ criteria fall on the same side of *k*_1_. As a result, the region between *k*_1_and the left-hand *k*_2_ criterion results in a low-confidence response despite being beyond the *k*_2_ boundary (relative to 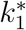). This occurs because this region is to the right of *k*_1_ and thus, due to asymmetric payoffs, the observer will make the *less* probable choice, which then results in low confidence in that choice. Effectively, the left-hand confidence criterion is shifted from *k*_2_ to *k*_1_. Here, we rely on the assumption that the confidence system is aware of the Type 1 decision (for further discussion of this issue, see Fleming and Daw, 2017).

The notion of an observer computing additional criteria for counterfactual reasoning is not new. For example, in the model of Type 1 conservatism of Maddox and Bohil (1998), where observers trade off gain versus accuracy, *k*_1_ is a weighted average of the optimal criteria for maximizing expected gain (*k*_*opt*_) and for exclusively maximizing accuracy (*k*_*p*_). In Zylberberg et al. (2018), observers learned prior probabilities of each stimulus type by an updating decision-making mechanism that computes the confidence the observer would have had if they had used the neutral criterion (*k*_*neu*_) for their Type 1 judgment. We suggest that for determining confidence in the face of asymmetric payoffs, normative observers compute the confidence they would have reported if they had instead used the *k*_*p*_ criterion for the discrimination judgment.

### 3.6 Type 2 Decision Models

In addition to the normative model we just described (i), we considered four sub-optimal models (ii-v) for the counterfactual Type 1 criterion about which the Type 2 criteria are yoked:

i. 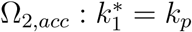
ii. 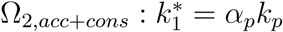
iii. 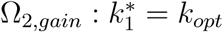
iv. 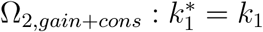
v. 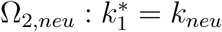

All of these models are characterized by the placement of the counterfactual criterion, 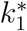; the distance *δ* is the only free parameter for all models and only alters the probability of a Type 2 response given the Type 1 response. That is, *δ* represents the propensity to respond low confidence, but the confidence criteria, *k*_2_, will be placed around 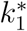 regardless of the particular value of *δ*. Thus, an observer’s overall confidence bias will be independent from a test of normativity. In the *normative-shift* model (Ω_2,*acc*_), the confidence criteria shift along with the discrimination criterion that maximizes accuracy and ignores possible payoffs. We also consider a *gains-shift* model in which confidence criteria shift with the criterion that maximizes expected gain (Ω_2,*gain*_), which is incorrect behavior in the case of asymmetric payoffs. In the *neutral-fixed* model (Ω_2,*neu*_), confidence criteria remain fixed around the neutral Type 1 criterion, regardless of the prior or payoff manipulation. Finally, for the classes of models that involve shifting confidence criteria (i.e., not the neutral-fixed model), we consider variants where conservatism in the discrimination criterion placement also affects 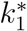: for the normative-shift model (Ω_2,*acc*+*cons*_) or the gains-shift model (Ω_2,*gain*+*cons*_). For the gains-shift model with carry-over conservatism, 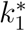 is identical to *k*_1_. For all other models, some combinations of priors and payoffs will decouple 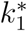 from *k*_1_. For the normative-shift model with carry-over conservatism, the decoupling only occurs for asymmetric payoffs. For the three remaining models, this decoupling occurs whenever priors and/or payoffs are asymmetric.

For simplicity, our models assume that the *k*_2_ criteria are placed symmetrically around 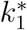 at a distance of ±*δ*. However, the ability to identify the underlying Type 2 model should not be affected by this assumption. Consider an observer whose low-confidence region to the left of 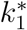 was always greater than their low-confidence region to the right of 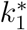, such that 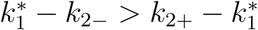. Then, the estimate of *δ* would be similar because the experimental design tested the mirror prior-payoff condition (i.e., for fixed *k*_2_, one condition would have 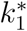 attracted to neutral and the other repelled, which is not the behaviour of 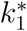 in any Type 2 model). Thus, the best-fitting model would be unlikely to change when *δ* is asymmetric, but the quality of the model fit would be impaired. Alternatively, an asymmetry in *δ* could be mirrored about the neutral criterion (e.g., the low confidence region closest to the neutral criterion is always smaller). Then, the *δ* asymmetry would be indistinguishable from a bias in the conservatism parameter. Although the confidence criteria are still yoked to 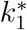, ultimately it is the patterns of confidence-criteria shift from all conditions jointly that are captured by the model comparison.

## 4 Methods

### 4.1 Participants

Ten participants (5 female, age range 22-43 years, mean 27.0 years) took part in the experiment. All participants had normal or corrected-to-normal vision, except one amblyopic participant. All participants were naive to the research question, except for three of the authors who participated. On completion of the study, participants received a cash bonus in the range of $0 to $20 based on performance. In accordance with the ethics requirements of the Institutional Review Board at New York University, participants received details of the experimental procedures and gave informed consent prior to the experiment.

### 4.2 Apparatus

Stimuli were presented on a gamma-corrected CRT monitor (Sony G400, 36 × 27 cm) with a 1280 × 1024 pixel resolution and an 85 Hz refresh rate. The experiment was conducted in a dimly lit room, using custom-written code in MATLAB version R2014b (The MathWorks, Natick, MA), with PsychToolbox version 3.0.11 (Brainard, 1997; Pelli, 1997; Kleiner et al., 2007). A chin-rest was used to stabilize the participant at a viewing distance of 57 cm. Responses were recorded on a standard computer keyboard.

### 4.3 Stimuli

Stimuli were Gabor patches, either right (clockwise) or left (counterclockwise) of vertical, presented on a mid-gray background at the center of the screen. The Gabor had a sinusoidal carrier with spatial frequency of 2 cycle/deg, a peak contrast of 10%, and a Gaussian envelope (SD: 0.5 deg). The phase of the carrier was randomized on each trial to minimize contrast adaptation.

### 4.4 Experimental Design

Orientation discrimination (Type 1, 2AFC, left/right) and confidence judgments (Type 2, 2AFC, low/high) were collected for seven conditions defined by the prior and payoff structure. The probability of a right-tilted Gabor could be 25, 50, or 75%. The points awarded for correctly identifying a rightversus a left-tilt could be 4:2, 3:3, or 2:4. In the 3:3 payoff scheme, a correct response was awarded 3 points. In the 2:4 and 4:2 schemes, correct responses were awarded 2 or 4 points depending on the stimulus orientation. Incorrect responses were not rewarded (0 points). We were interested in people’s natural confidence behavior, so confidence responses were not rewarded, allowing participants to respond with their subjective sense of probability correct. The prior and payoff structure was explicitly conveyed to the participant before the session began (Fig. 2b) and after every 50 trials. There were 7 prior-payoff conditions (Fig. 2c): no asymmetry (50%, 3:3), single asymmetry (50%, 4:2; 50%, 2:4; 25%, 3:3; 75%, 3:3), or double asymmetry (25%, 4:2; 75%, 2:4). Note that two of the possible double asymmetry conditions (25%, 2:4; and 75%, 4:2) were not tested because these conditions incentivized one response alternative to such a degree that they would not be informative for model comparison. Participants first completed the full-symmetry condition, followed by the single-asymmetry conditions in random order, and finally the double-asymmetry conditions, also in random order (Fig. 2d). Session order facilitated task completion and participants’ understanding of the prior and payoff asymmetries before encountering both simultaneously. Each condition was tested in a separate session with no more than one session per day. In all sessions, participants were instructed to report their confidence in the correctness of their discrimination judgment.

**Figure 2:**
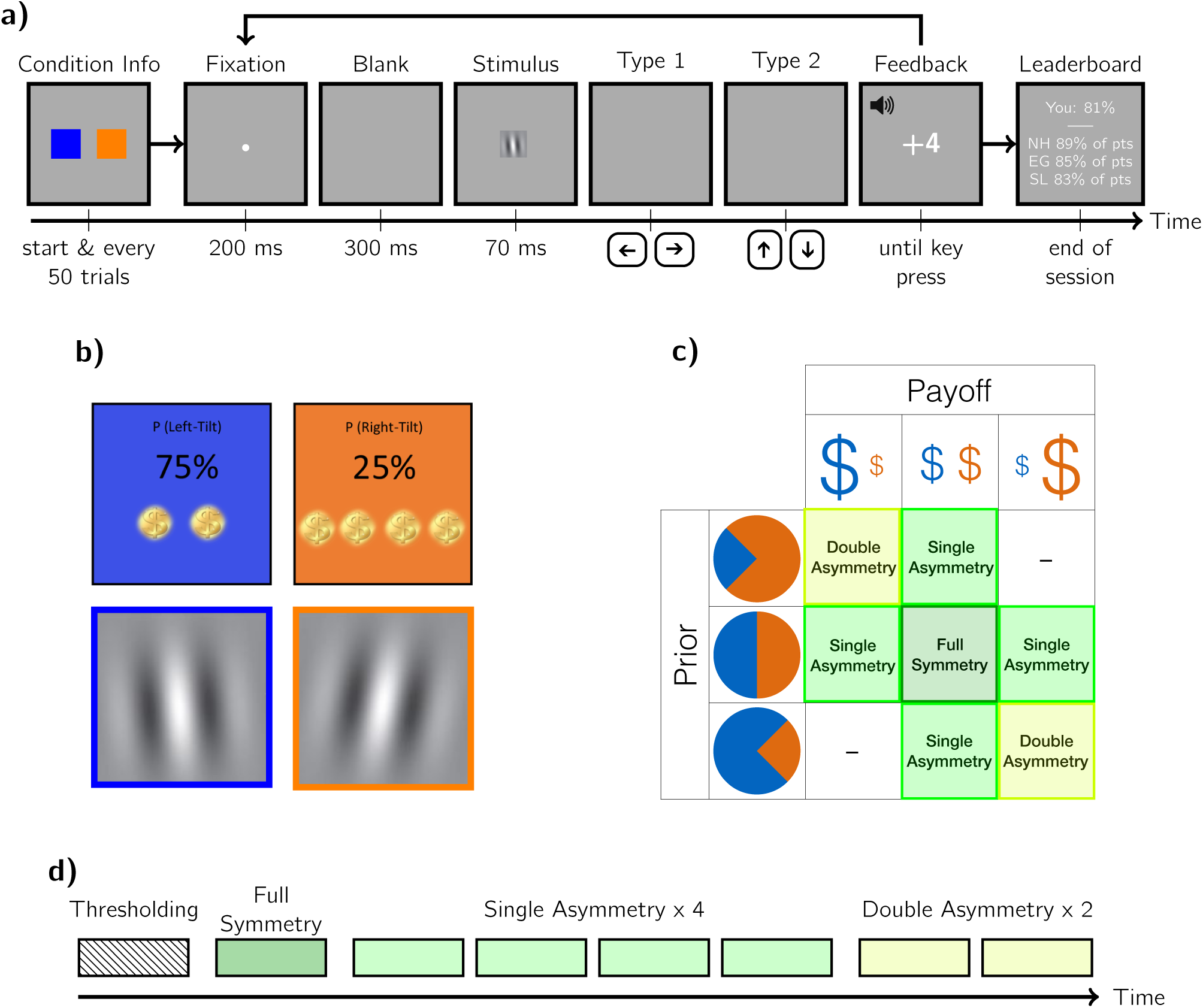
Experimental methods. a) Trial sequence including an outline of the initial condition information screen (see part (b) for details) and final (mock) leaderboard screen. Participants were shown either a right- or left-tilted Gabor and made subsequent Type 1 and Type 2 decisions before being awarded points and given auditory feedback based on the Type 1 discrimination judgment. b) Sample condition-information displays from a double-asymmetry condition. Below: Example Gabor stimuli, color-coded blue for left- and orange for right-tilted. The exact stimulus orientations depended on the participant’s sensitivity. c) Condition matrix. Pie charts show the probability of stimulus alternatives (25, 50, or 75%) and dollar symbols represent the payoffs for each alternative (2, 3, or 4 pts). Squares are colored and labeled by the type of symmetry. d) Timeline of the eight sessions. The order of conditions was randomized within the single- and within the double-asymmetry conditions.

### 4.5 Thresholding Procedure

A thresholding procedure was performed prior to the main experiment to equate difficulty across observers to approximately *d*′ = 1. Observers performed a similar orientation-discrimination judgment as in the main experiment. Absolute tilt magnitude varied in a series of interleaved 1-up-2-down staircases to converge on 71% correct. Each block consisted of three staircases with 60 trials each. Participants performed multiple blocks until it was determined that performance had plateaued (i.e., learning had stopped). Preliminary thresholds were calculated using the last 10 trials of each staircase. At the end of each block, if none of the three preliminary thresholds were better than the best of the previous block’s preliminary thresholds, then the stopping rule was met. As a result, participants completed a minimum of two blocks and no participant completed more than five blocks. A cumulative Gaussian psychometric function was fit by maximum likelihood to all trials from the final two blocks (360 trials total). The slope parameter was used to calculate the orientation corresponding to 69% correct for an unbiased observer (*d*′ = 1; Macmillan and Creelman, 2005). This orientation was then used for this subject in the main experiment. Thresholds ranged from 0.36 to 0.78 deg, with a mean of 0.59 deg.

### 4.6 Main Experiment

Participants completed seven sessions, each of which had 700 trials with the first 100 treated as warm-up and discarded from the analysis. All subjects were instructed to hone their response strategy in the first 50 trials to encourage stable criterion placement. The trial sequence is outlined in Fig. 2a. Each trial began with the presentation of a fixation dot for 200 ms. After a 300 ms inter-stimulus interval, a Gabor stimulus was displayed for 70 ms. Participants judged the orientation (left/right) and then indicated their confidence in that orientation judgment (high/low). Feedback on the orientation judgment was provided at the end of the trial by both an auditory tone and the awarding of points based on the session’s payoff structure. Additionally, the running percentage of potential points earned was shown on a leaderboard at the end of each session to foster inter-subject competition. Participants’ cash bonus was calculated by selecting one trial at random from each session and awarding the winnings from that trial, with a conversion of 1 point to $1, capped at $20 over the sessions. Total testing time per subject was approximately 8 hrs.

### 4.7 Model Fitting

Detailed description of the model-fitting procedure can be found in the Supplementary Information (Sections 1 and 2). Briefly, model fitting was performed in three sequential steps. First, we estimated a per-participant *d*′ and meta-*d*′ using a hierarchical Bayesian model. We used as inputs the empirical *d*′ and meta-*d*′ calculated separately for each prior-payoff condition. These per-participant sensitivities were fixed for all subsequent modeling. Second, we fit the discrimination behaviour according to the Type 1 models, selecting the best-fitting Type 1 model for each participant before the final step of fitting the confidence behavior according to the Type 2 models. For the Type 1 and Type 2 models, we calculated the log likelihood of the data given a dense grid of parameters (*α, γ*, and *δ*) using multinomial distributions defined by the stimulus type, discrimination response, and confidence response. All seven prior-payoff conditions were fit jointly. Model evidence was calculated by marginalizing over all parameter dimensions and then normalizing to account for grid spacing.

## 5 Results

We sought to understand how observers make perceptual decisions and confidence judgments in the face of asymmetric priors and payoffs. Participants performed an orientation-discrimination task followed by a confidence judgment. To account for the behavior, we defined two sets of models. Type 1 models defined the contribution of conservatism to the discrimination responses. Type 2 models defined the role of priors, payoffs, and conservatism in the confidence reports. We were interested in which of three classes of models best fits confidence behavior: neutral-fixed, gains-shift, or normative-shift.

### 5.1 Model Fits

Type 1 models were first fit using the discrimination responses alone. Four models were compared: optimal criterion placement (Ω_1,*opt*_), equal conservatism for priors and payoffs (Ω_1,1*α*_), different degrees of conservatism for priors and payoffs (Ω_1,2*α*_), and a model in which there was a failure of summation of criterion shifts in the double-asymmetry condition (Ω_1,3*α*_). Fitting the Type 1 models also provided an estimate of left/right response bias, *γ*. We performed a Bayesian model selection procedure using the SPM12 Toolbox (Wellcome Trust Centre for Neuroimaging, London, UK) to calculate the protected exceedance probabilities (PEPs) for each model (Figure 3a). The exceedance probability (EP) is the probability that a particular model is more frequent in the general population than any of the other tested models. The PEP is a conservative measure of model frequency that takes into account the overall ability to reject the null hypothesis that all models are equally likely in the population (Stephan et al., 2009; Rigoux et al., 2014). Overall, an additional parameter in the double-asymmetry conditions was needed to explain Type 1 criterion placement, indicating a failure of summation of criterion shifts (i.e., the best-fitting model was Ω_1,3*α*_).

**Figure 3:**
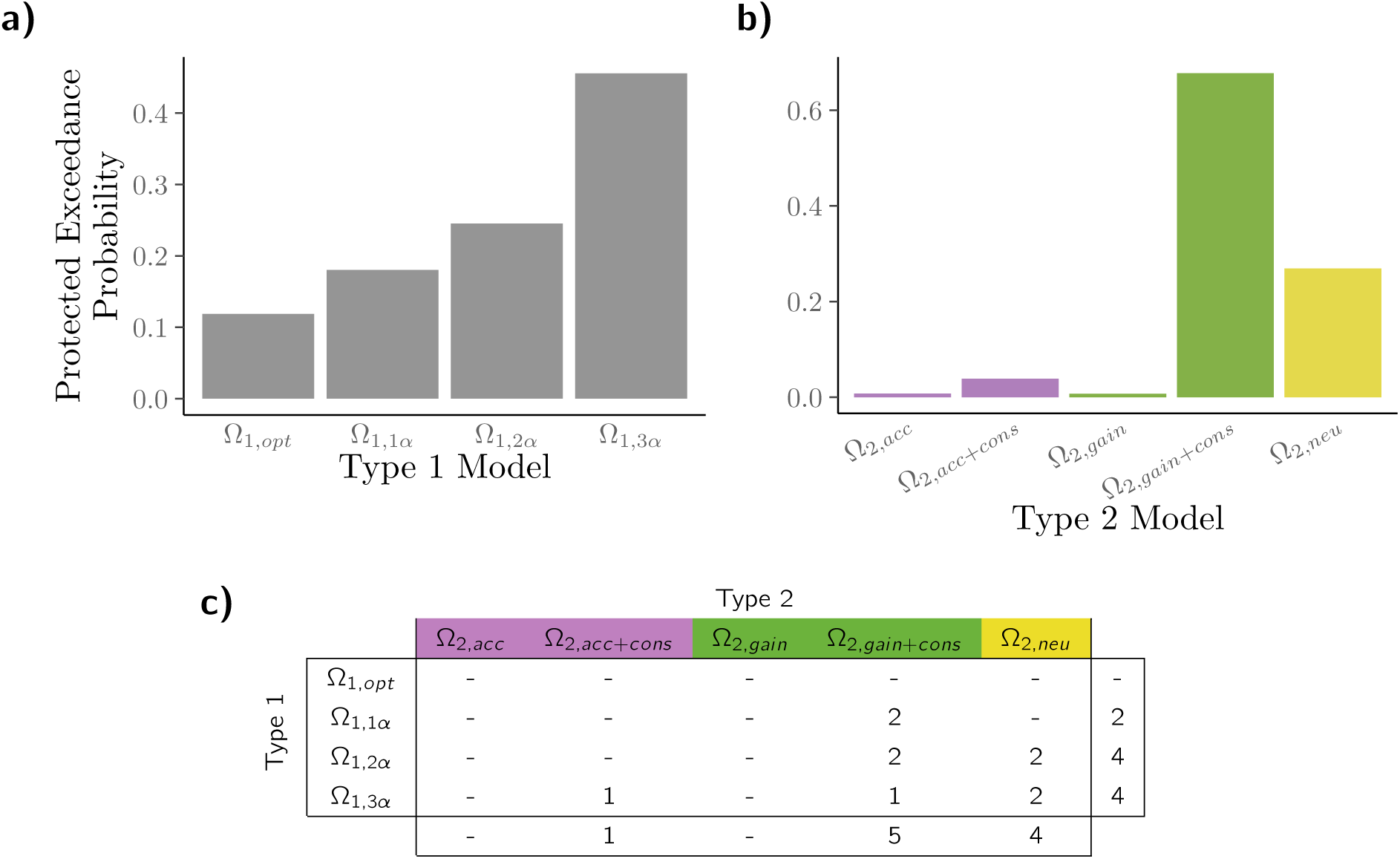
Model comparison for the Type 1 and Type 2 responses. a) The protected exceedance probabilities (PEPs; see text for details) of the four Type 1 models. b) PEPs of the five Type 2 models. Note that model comparisons were performed first for Type 1 and then for Type 2 responses, using the best-fitting Type 1 model and parameters, on a per-subject basis, in the Type 2 model evaluation. c) Best-fitting models for each participant. Purple: normative-shift models, green: gains-shift models, yellow: neutral-fixed model.

In the second step, the Type 2 models were fit using each participant’s best Type 1 model and the associated maximum *a posteriori* (MAP) parameter estimates. The Type 2 models differed in the placement of the Type 2 criteria, which split the internal response axis into “high” and “low” confidence regions, for each “right” and “left” discrimination response. We modeled the two Type 2 criteria as shifting to account for only the prior probability, maximizing accuracy with the confidence response (Ω_2,*acc*_; normative-shift class), shifting the confidence criteria in response to payoff manipulations (Ω_2,*gain*_; gains-shift class), or failing to move the confidence criteria away from neutral at all (Ω_2,*neu*_; neutral-fixed class). For the models with shifted confidence criteria, we also tested for effects of Type 1 conservatism on Type 2 decision-making (Ω_2,*acc*+*cons*_ and Ω_2,*gain*+*cons*_; both sub-optimal). We again compared the models quantitatively with PEPs (Figure 3b). The favored model was the gains-shift model with carry-over conservatism, Ω_2,*gain*+*cons*_. This model shifts the confidence criteria in response to both prior and payoff manipulations with the conservatism that participants exhibited in the Type 1 decisions affecting placement of the confidence criteria.

Figure 3c shows the best-fitting models for individual participants, according to the amount of relative model evidence (here the marginal log-likelihood). All of the sub-optimal Type 1 models (i.e., not Ω_1,*opt*_) were a best-fitting model for at least one of the ten participants. Similarly, no one was best fit by the normative-shift without Type 1 conservatism either (Ω_2,*acc*_). Overall, there was no clear pattern between the pairings of Type 1 and Type 2 models.

### 5.2 Model Checks

We performed several checks on the fitted data to ensure that parameters were capturing expected behavior and that the models could predict the data well (reported in detail in Section 3 of the Supplementary Information). The quality of a model is not only dependent on how much more likely it is than others, but it is also dependent on its overall predictive ability. To visualize each model’s ability to predict the proportion of each response type (“right” vs. “left” x “high” vs. “low”), we calculated the expected proportion of each response type given the MAP parameters for each model and participant. We compared the predicted response proportions to the empirical proportions (Figure 4). Larger residuals are represented by more saturated colors. For the best-fitting models, the residuals are small and unpatterned.

**Figure 4:**
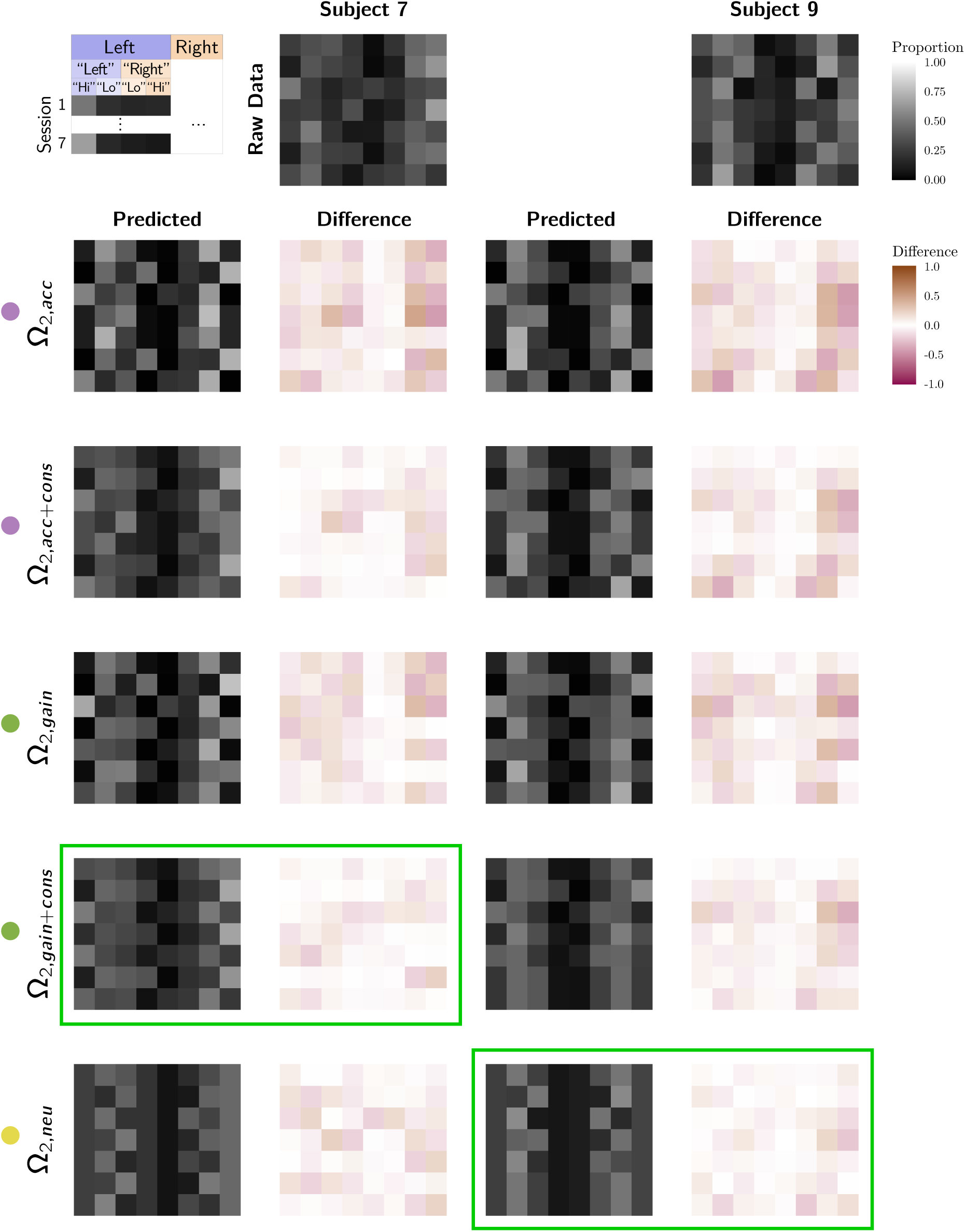
Visualization of the raw and predicted response rates for two example participants. Grids are formed of the seven conditions (rows) and the eight possible stimulus-response-confidence combinations (columns). See Figure S3 in the Supplement for condition order. The fill indicates the proportion of trials for that condition and stimulus that have that combination of response and confidence. Top row: Raw response rates of two example subjects. Subsequent rows, columns 1 and 3: Predicted response rates for each Type 2 model using the best-fitting parameters of the best-fitting Type 1 model for that individual. Columns 2 and 4: Difference between raw and predicted response rates. Green boxes: winning models (Subject 7: Ω_*gain*+*cons*_; subject 9: Ω_*neu*_). Colored circles by model names indicate purple: normative-shift models, green: gains-shift models, yellow: neutral-fixed model.

We also compared the Type 1 criteria and the counterfactual confidence criteria (Figure 5). We constrained the empirical counterfactual confidence criterion to be the midpoint between the two Type 2 criteria (i.e., 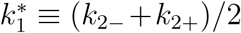). Using 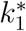, the predictions made by the Type 2 models are highly distinguishable. In the left-most column, predicted *k*_1_ and 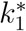 for each session are shown for each model, assuming *d*′ = 1 and either Ω_1,*opt*_ or Ω_1,1*α*_ where *α* = 0.5. In the top row, empirical criteria from the same two example participants as in Figure 4 are shown. Empirical criteria are calculated with the standard SDT method (detailed in Section 1 of the Supplementary Information, see Figure S1).

**Figure 5:**
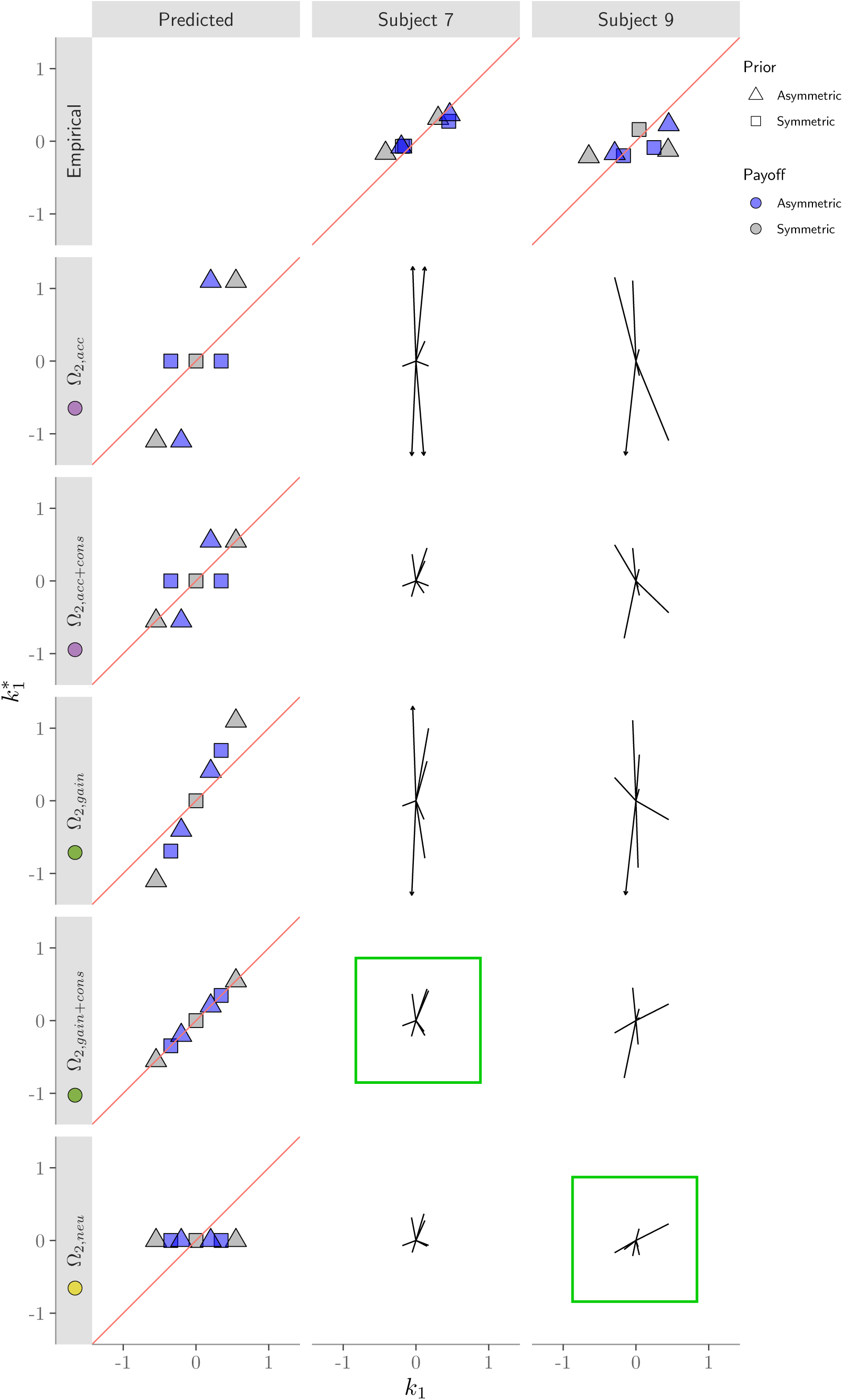
Comparison of the empirical and predicted *k*_1_ and 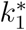. Top row: empirical criteria of two example observers. The 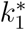 was calculated as the midpoint between the two empirical *k*_2_ (see Figure S1 for *k*_2_ calculation details). Left column: predicted relationship between the Type 1 and Type 2 criteria (*d*′ = 1; all Ω_1,1*α*_ with *α* = 0.5). Grey and square symbols: symmetry conditions. Triangles: prior asymmetry. Blue symbols: payoff asymmetry. Polar plots: residuals between empirical data and model prediction based on best-fitting parameters, plotted as vectors. Arrowheads: residuals greater than plot bounds. Colored circles by model names indicate purple: normative-shift models, green: gains-shift models, yellow: neutral-fixed model.

The visualization in the top row and left-most column of Figure 5 illustrates several behavioral phenomena. The response bias, *γ*, results in a shift in all criteria in the same direction, translating all data points parallel to the identity line. Conservatism is represented by an attraction of all data toward the origin on the x-axis for Type 1 and the y-axis for Type 2 judgments. The Type 2 models predict qualitatively different arrangements of the data points. If the prior and payoff asymmetries affect the placement of the Type 1 criterion but not the Type 2 criteria (Ω_2,*neu*_; neutral-fixed), the data are clustered along a single value on the y-axis. If the prior and the payoff affect the placement of the Type 1 and Type 2 criteria equally, (Ω_2,*gain*_; gains-shift), then the data fall on the identity line. With normative-shift behavior (Ω_2,*acc*_), the prior asymmetry conditions (grey triangles) fall on the identity line because confidence tracks the prior, while in the payoff asymmetry conditions (blue squares), the data have the same *k*_2_ midpoint as in the neutral condition (grey squares) because confidence does not track the payoff.

Vectors in all 10 of the bottom right polar plots represent the difference (i.e., the residual) between the empirical and the predicted criteria from the model fits. While the model prediction column is based on fixed parameters, the predicted data in the 10 polar plots use parameters that best fit the participant’s data using that model. It is immediately clear that the normative-shift model without carry-over conservatism (second row) does a poor job of describing participants’ behavior, and that, in general, conservatism is a necessary component of both the Type 1 and Type 2 models.

### 5.3 Type 1 Conservatism

While not the main focus of the study, it was important to consider the role of Type 1 conservatism to properly capture the Type 1 decision-making behavior. First, we remark on the relative magnitude of conservatism due to priors and payoffs. Figure 6a shows fitted *α*_*p*_ and *α*_*v*_ under the most complex conservatism model (Ω_1,3*α*_) and Figure 6b shows them under the best-fitting model for each observer. These figures show that eight of the ten participants were conservative in their criterion placement for both prior and payoff manipulations, as indicated by data points in the gray regions. Of the eight participants that displayed conservatism, five were significantly more conservative for payoff asymmetries than prior asymmetries (*α*_*v*_ < *α*_*p*_), whereas only one was significant in the opposite direction (*α*_*p*_ < *α*_*v*_). At the group level, however, we did not find a significant difference between the best fitting *α*_*v*_ and *α*_*p*_, either for the best-fitting Type 1 model or the winning model (paired *t*-tests, *p* > 0.05). Note that the negative *α* values derive from a participant who shifted criteria consistently in the opposite direction expected from a rational observer in response to manipulations of payoffs and priors.

**Figure 6:**
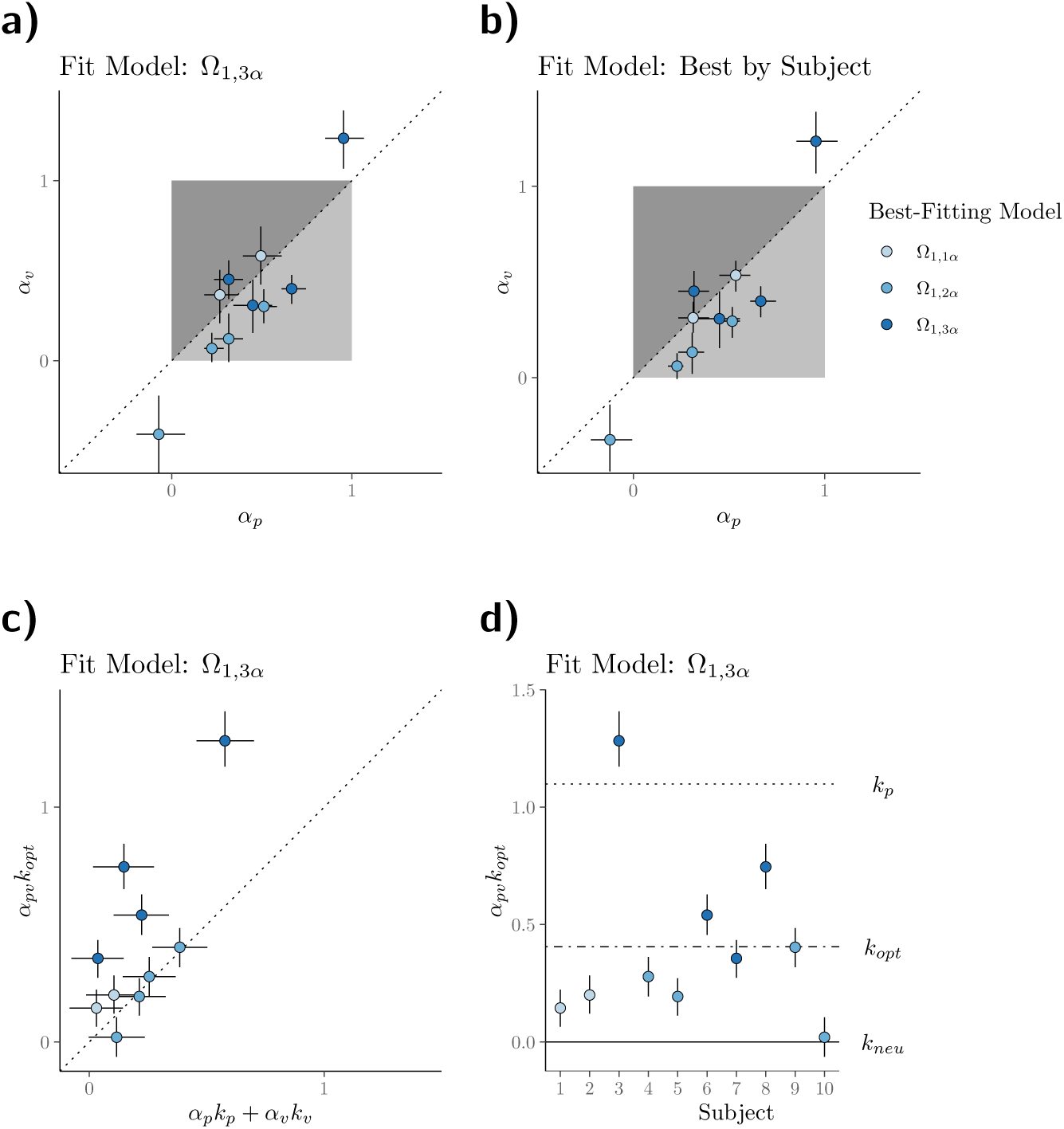
Conservatism for Type 1 decision-making. a) A comparison of the extent of conservatism under payoff versus prior asymmetries. Each data point represents the best-fitting conservatism parameters of a single observer when fit by Ω_1,3*α*_. These parameters are only contingent on the conservatism in the single-asymmetry conditions. In this model, conservatism in the double-asymmetry conditions is captured by a separate model parameter. Darker marker fill: additional conservatism parameters were required to fit to that observer’s data. Dashed line: equality line. Dark grey region: conservatism greater for prior than payoff manipulations (i.e., *α*_*p*_ < *α*_*v*_). Light grey region: conservatism is greater for payoffs (i.e., *α*_*p*_ > *α*_*v*_). Data points outside these regions are not consistent with conservative criterion placement. b) Same as (a) using fit parameters from the best-fitting Type 1 model for each observer. c) Test of summation of criterion shifts using the Ω_1,3*α*_ model fits. Observers who required a third *α* to capture their data (i.e., were best fit by Ω_1,3*α*_) had criterion shifts for the double-asymmetry conditions that were not well predicted as the sum of the shifts in the single-asymmetry conditions. d) Criterion placement in the double-asymmetry conditions. These are the same data as in the y-axis of (c), but extended to more easily compare the actual criterion placement with potential other task-relevant criteria. Horizontal criteria lines assume *d*′=1.

An additional implication of SDT is that an ideal observer’s criterion shift due to payoffs and due to priors should sum (Stevenson et al., 1990): *k*_*pv*_ = *k*_*p*_ + *k*_*v*_ (Figure 1b). Figure 6c contrasts the prediction of this additive rule with the empirical results. The difference between the predicted and actual criterion shift is significant (*t* = 2.41, *p* =.039), with the effect primarily driven by the four observers best fit by the non-additive conservatism model, Ω_1,3*α*_. Each of these four observers had 95% CIs that did not overlap with the identity line. We show the criterion placement in the double-asymmetry cases in Figure 6d. Most observers did not shift their criterion far enough from neutral to the optimal placement, *k*_*opt*_. Three observers, however, placed their criterion beyond *k*_*opt*_, with two stopping short of the accuracy-maximizing criterion *k*_*p*_.

## 6 Discussion

### 6.1 Confidence Behavior

The primary focus of this study was to assess how observers assigned confidence to the discrimination decision for different prior-payoff scenarios. Three Type 2 model classes were characterized by the placement rule for the counterfactual Type 1 criterion, 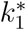, to which the confidence criteria, *k*_2_, were yoked. The classes were defined by the counterfactual criterion coinciding with the accuracy-maximizing criterion (normative-shift), the gain-maximizing criterion (gains-shift), or the neutral criterion (neutral-fixed). The majority of observers were best explained by the gains-shift model with carry-over conservatism (Ω_2,*gain*+*cons*_) or the neutral-fixed model (Ω_2,*neu*_), with the Bayesian model selection favoring the former. One participant was best fit by the normative-shift model with carry-over conservatism (Ω_2,*acc*+*cons*_). Furthermore, we found no clear pattern between the number of Type 1 conservatism parameters required to explain discrimination behavior and the placement strategy for confidence criteria.

For the subset of observers who were best fit by the neutral-fixed model, the perceived tilt magnitude was predictive of confidence in all prior-payoff scenarios. While these observers correctly did not allow the payoff structure of the environment to affect confidence, it was non-normative to ignore the additional information provided by the priors for the response alternatives. This notion of ‘sticky’ or fixed confidence criteria has been examined previously in the context of changing stimulus reliability, where confidence criteria should be shifted to avoid a preponderance of high-confidence reports for the low-reliability stimuli. Empirical results are mixed; Zylberberg et al. (2014) found participants were reluctant to shift their criteria sufficiently to account for the different reliabilities, whereas a fixed-criterion model was rejected by Adler and Ma (2018). Our results suggest that some observers can be insensitive to the prior-payoff context when it comes to placing confidence criteria, despite our efforts to present each prior-payoff context in separate sessions, keep stimulus reliability and attentional factors constant, and provide substantial context information and training.

In contrast, the confidence criteria of gains-shift observers tracked the placement of the criterion used for the Type 1 judgment. As such, priors were correctly incorporated into confidence judgments but payoffs were inappropriately incorporated also. For such people, higher relative reward leads to selection of the highly rewarded alternative and, on average, higher confidence about reporting that outcome. In effect, gains-shift behavior can be viewed as a naïve optimism for selecting the highly rewarded outcome: “this highly rewarding perceptual alternative that I have selected is certainly the state of the world”. This bias for higher confidence with greater reward is consistent with what has been reported previously in the perceptual lottery tasks of Lebreton et al. (2018).

The finding that most observers did not appropriately dissociate Type 1 and Type 2 criteria is compelling, particularly so in the case of the gains-shift observers. By not selectively decoupling their *k*_1_ and 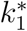 for asymmetric payoffs, these observers faced a trade-off between maximizing gains with the discrimination report and faithfully representing perceived accuracy with the confidence report. Consider the following real-world example of a pilot judging whether their aircraft is heading for collision with an upcoming mountain peak using weak sensory evidence (e.g., night time or fog). A normative-shift pilot would make a corrective action because of the high cost of collision, but not be confident that a collision will occur. In contrast, a gains-shift pilot would similarly adjust the aircraft heading, but would also be likely to have high confidence that the collision was imminent despite the weak sensory evidence. The experience of the gains-shift pilot in a world full of dangerous possibilities would be unsettling. However, if this gains-shift pilot places their discrimination criterion somewhere between the gains-maximizing criterion and the accuracy-maximizing criterion (i.e., payoff conservatism), then their confidence judgments will better reflect the true state of the world. Subsequent laboratory experiments can examine this trade-off by using more complex reward structures and/or elaborated decision scenarios.

We also note some simple experimental factors that may have produced the observed pattern of confidence results. First, the lack of adaptability of the neutral-fixed observers should not be taken as evidence of an inability to adapt. It is possible that these observers ignored the prior-payoff structure entirely for confidence because it changed from session to session, and instead opted for a criterion-placement strategy that would work best for all conditions of the experiment. This is unlikely, however, because they did not adopt such a strategy for discrimination. For the gains-shift observers, we note that a failure to understand the task instructions could explain their behavior. It is possible that observers did not report the probability they were correct, as per the experimenter instructions, but instead considered their expected gain from the trial when reporting confidence. However, all participants deviated from normative behavior, making it unlikely that these experimental factors alone can explain our results.

### 6.2 Discrimination Behavior

Observers were generally conservative in the placement of the discrimination criterion, *k*_1_, as most participants were best described by a model with some form of conservatism, with the majority best fit with two or three separate *α* parameters. In the Type 1 model comparison, the winner was the non-additive conservative model (Ω_1,3*α*_), where three *α* parameters were needed to capture discrimination behaviour (Healy and Kubovy, 1981). Despite Bayesian model selection favoring the non-additivity model, only 40% of our sample population was best fit by this model, which is as similarly inconclusive as it was for previous attempts at testing additivity (Stevenson et al., 1990). We found significant differences at the individual-subject level, but not at the group level, that conservatism was stronger when the payoffs were asymmetric than when the priors were asymmetric. Thus, the observed differences in conservatism for priors and payoffs in our study were less apparent than in most previous studies (Lee and Zentall, 1966; Ulehla, 1966; Healy and Kubovy, 1981; Maddox, 2002; Ackermann and Landy, 2015), but not all (Healy and Kubovy, 1978).

Several factors may have contributed to the observed Type 1 conservatism. One hypothesis is that observers trade off between maximizing gains and maximizing accuracy (Maddox and Bohil, 1998), as it may be hard for the observer to sacrifice accuracy for expected gain. In Section 4 of the Supplementary Information, we demonstrate that gain-accuracy trade-off model of conservatism is equivalent to our Ω_1,2*α*_ model, indicating that the gain-accuracy trade-off strategy alone cannot account for the observed non-additivity. Alternatively, conservatism could depend on the criterion-adjustment strategy (Busemeyer and Myung, 1992), which suggests that observers will not shift their criterion far from neutral for an inconsequential gain, causing them to fall short of optimal. Non-additivity is possible due to the non-linear effects on the slope of the expected-gain function from combining asymmetric priors and payoffs. However, 30% of observers placed their criterion beyond the optimal criterion in the double-asymmetry conditions, which is inconsistent with a reluctance to shift the criterion sufficiently from neutral. In fact, these criteria are biased in the direction of the accuracy-maximizing criterion, as would be expected under the gain-accuracy trade-off hypothesis. A mix of gain-accuracy trade-off strategy and criterion-adjustment strategy (Maddox and Bohil, 2003), that could produce both unequal conservatism and non-additivity, would better explain our results.

A metacognitive source of conservatism proposed by Kubovy (1977) implicates *d*′ in Eq. 5. Observers likely form an estimate of their overall performance from experience with the task. If they happen to overestimate performance (i.e., 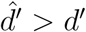), then it follows from Eq. 5 that *k*_1_ < *k*_*opt*_. Note that this is not confidence for a given discrimination response, but a metacognitive appraisal of the difficulty of the task, such as the expected performance indicated by the uncertainty in the stimulus (Zylberberg et al., 2014). According to this hypothesis, most of the observers would have been overestimating performance to be conservative, with the one observer with liberal criterion placement underestimating their performance. Observations of overconfidence are a common finding in metacognitive studies (Baldassi et al., 2006; Mamassian, 2008; Zylberberg et al., 2014; Mamassian, 2016; Lebreton et al., 2018; Charles et al., 2020) as is conservatism (Lee and Zentall, 1966; Ulehla, 1966; Healy and Kubovy, 1981; Maddox, 2002; Ackermann and Landy, 2015). However, overestimation of discrimination performance by itself is an insufficient explanation for conservatism, as it cannot explain the differences in the degree of conservatism for priors versus payoffs, observed for some participants, or the non-additivity results. However, if performance estimations differ under manipulations of priors versus payoffs, specifically larger overestimations of performance for asymmetric payoffs, conservatism would be larger for payoffs than priors. Furthermore, it is entirely plausible that the contribution of priors and payoffs to performance estimation is non-linear, which would result in non-additivity of criteria.

Finally, we note that a simple experimental factor may have encouraged conservatism in general. By starting testing with the symmetrical prior-payoff design in the thresholding procedure and initial testing session, this session order may have encouraged participants to anchor the Type 1 decisions to the neutral criterion. However, this explanation cannot account for observed unequal conservatism or non-additivity. Overall, we conclude that the conservatism observed in this task is likely due to more than one of the following possible factors: noisy behavior, strategies to trade off gain versus accuracy, sub-optimal criterion adjustment, and biases in participants’ judgments of their own *d*′.

### 6.3 Type 1 Conservatism Applied to Type 2 Judgments

It is currently a matter of debate whether the internal sensory measurement used by the perceptual decision-making system is the same or similar to that used by the metacognitive decision-making system (e.g., Resulaj et al., 2009; Fleming and Daw, 2017; Peters et al., 2017). The standard SDT framework assumes the same internal measurement is used for both Type 1 and 2 judgments. However, there is substantial evidence to suggest that additional noise is applied to the internal measurement between the Type 1 and 2 judgments (Maniscalco and Lau, 2012; Fleming and Lau, 2014; Maniscalco and Lau, 2016; Bang et al., 2019). We found supporting evidence of additional metacognitive noise in the form of reduced metacognitive sensitivity (a ratio of meta-*d*′ to *d*′ of 0.86 ± 0.04; see Section 1 of the Supplementary Information), which we incorporated into our SDT model. We also consistently found that Type 1 conservatism carried over into the Type 2 confidence-criteria placement for the observers best fit by the gains-shift and normative-shift model classes. This raises a different, but related question: to what extent are decision-related parameters of the system, such as criteria placement, shared between the perceptual and metacognitive systems? And how is this information shared from the Type 1 to the Type 2 response? We speculate on several possibilities.

First is the simplest scenario: the Type 1 and Type 2 processes are computed jointly using the same information, with confidence being an additional readout of the same decision mechanism. However, in addition to the evidence of additional metacognitive noise, there is considerable evidence that neural processing occurs in distinct regions for perceptual and metacognitive decision-making (Shimamura, 2000; Fleming and Dolan, 2012; Rahnev et al., 2016; Shekhar and Rahnev, 2018), suggesting this is unlikely the case.

Second, the Type 1 system might convey only relative information to the Type 2 system, such as how far the measurement was from the decision boundary, rather than noisily propagating the internal measurement itself. In this scenario, the additional metacognitive noise could be a result of computing this difference. A relative measurement also has the advantage over the other hypotheses that it only requires one piece of information to be sent to the Type 2 system (i.e., the relative measurement and not context information). Despite being efficient, this hypothesis is not supported by our results. Given that the neutral-fixed observers were able to dissociate *k*_1_ and 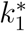 by keeping the latter fixed at the neutral criterion, this suggests that the Type 2 system does not receive an internal measurement coded relative to the discrimination criterion *k*_1_.

Third, the Type 2 system might be independent of the Type 1 system, but receives the same context information. It also produces conservatism, but is flexible enough to allow 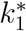 to be independent of *k*_1_. All types of observers (gains-shift, normative-shift, and neutral-fixed) can be explained by such a Type 2 system. However, given the flexibility of such a system, why weren’t observers able to reduce the influence of the payoff structure at the second processing step?

Fourth, the Type 1 system might directly inform the Type 2 system of both its decision boundary *k*_1_ and the internal measurement. The gains-shift observers then can yoke their confidence criteria to *k*_1_, whereas the neutral-fixed observers can ignore this extra *k*_1_ signal. This system is both relatively simple and explains the results of the majority of observers. Thus, we favor this interpretation, in which the discrimination decision boundary is propagated to the Type 2 system. Further work is required to understand why normative metacognitive behavior was not achieved, and why some observers may or may not incorporate *k*_1_ into their confidence judgment.

### 6.4 Criterion Stability

Often of concern when conducting psychophysical experiments is whether assumptions of criterion stability are valid. In some circumstances, criterion shifts within an experimental session are appropriate and expected, such as Type 1 criterion shifts in response to variable priors (Norton et al., 2017; Zylberberg et al., 2018) or Type 2 criterion shifts when intermixing stimuli of different difficulties (Zylberberg et al., 2014; Adler and Ma, 2018). In scenarios where they should be fixed, the best practices to encourage stable criteria are to only use one pair of stimuli (i.e., fixed difficulty), collect the data in a single session, and to not combine data across participants (Macmillan and Creelman, 2005). We met all recommendations, as we ensured context effects were kept constant within an individual session as would be done for fixing difficulty (although our models did include some assumptions about criterion stability across sessions, see Models). However, issues of unstable criteria can occur even in studies with unchanging contexts (e.g., Yu and Cohen, 2009) or fixed difficulty (e.g., Maniscalco and Lau, 2012). Type 2 criterion instability, indicated in the latter example, is mathematically equivalent to additional noise between the perceptual and confidence decisions (Maniscalco and Lau, 2016), which our models incorporate as meta-*d*′ (see Supplementary Information, Section 1). But, more generally, how may criterion instability interact with our models? No particular patterns were evident between the best-fitting model and estimated *d*′, meta-*d*′, or their ratio (Supplementary Information, Section 1), although a larger sample of participants is likely needed to resolve any small differences. Otherwise, we predict that unstable criteria impact the overall quality of the model fits, but do not introduce a bias in the Type 2 model selection.

### 6.5 Conclusion

By manipulating priors and payoffs in a perceptual task, we found sub-optimal decision-making at the Type 1 level and non-normative decision-making at the Type 2 level. Discrimination judgments were conservative, with similar conservatism for payoffs and priors, and non-additivity of criterion shifts when both priors and payoffs were asymmetric. Confidence judgments were non-normative in one of two ways: 1) observers did not consider the role of priors or 2) they incorporated payoffs, which accord with the neutral-fixed and gains-shift classes of models respectively. Both of these strategies hinder decision-making. For example, a radiologist who ignores prior probabilities when assigning confidence might hesitate to recommend further tests for a patient who is a heavy smoker. Similarly, a radiologist who inappropriately incorporates payoffs may be more confident in a positive diagnosis if he receives kickbacks from the imaging center to encourage future scans. The patterns of behavior found in this task point to explanations of why humans may consider trade-offs between maximizing gain and maximizing accuracy, as well as provide new insights about the role of the decision boundary in Type 1 versus Type 2 computations.

## Acknowledgements

This work was supported by NIH grant EY08266, NIH Training Program in Computational Neuroscience grant T90DA043219, NSF Collaborative Research in Computational Neuroscience grant 1420262, NSF GRFP DGE1342536, and French ANR grant ANR-18-CE28-0015-01 “VICONTE”.

## Author Note

This research was presented at the 2018 Vision Sciences Society meeting at St. Pete Beach, Florida. It has also been made available in manuscript form on the BioRxiv server at https://www.biorxiv.org/content/10.1101/703082v1. Data from this study can be found at https://osf.io/d6ef3/. This study was not pre-registered. Individual author contributions in the CRediT taxonomy style are as follows (black: major contribution, gray: minor contribution, see https://www.casrai.org/credit.html for more details):

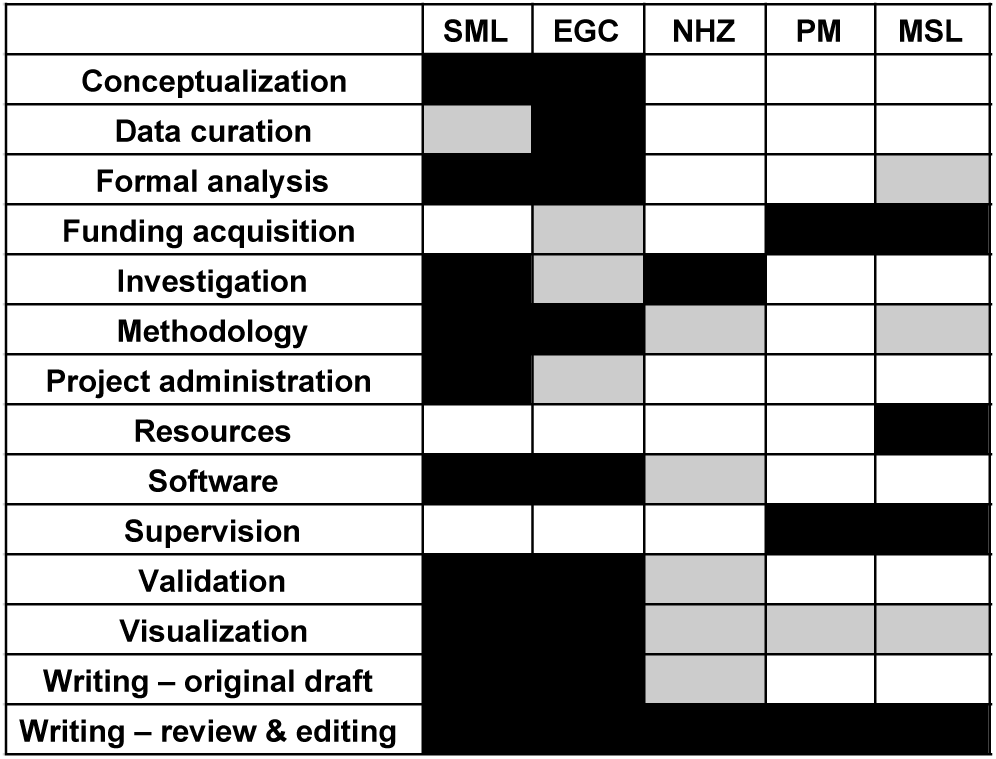

