## Supplementary Information for "Priors and Payoffs in Confidence Judgments"

### Contents

**1 Type 1 and Type 2 Sensitivity** **1**

**2 Multinomial Decision Model** **5**

**3 Model Checks and Fits for All Subjects** **7**

**4 Gains-Accuracy Trade-off Strategy and Conservatism** **13**

### 1 Type 1 and Type 2 Sensitivity

To fit the models presented in this paper, we required an estimate of discrimination sensitivity ($d'$) and metacognitive sensitivity (meta-$d'$) for each observer. Each participant completed a threshold procedure to find the Gabor orientation that would yield a $d'$ of 1. We could have used this for all analyses, however we sought to utilize all of the decisions made in the main task to better estimate $d'$, as well as obtain a reasonable estimate of meta-$d'$. To achieve this, we implemented a hierarchical Bayesian model that leveraged all possible sources of information to yield a single estimate of $d'$ and meta-$d'$ for each participant. We computed the empirical $d'$ for participant $i$ in session $j$ of the main task according to the standard formula

$$d'_{ij} = z(p\text{H}_{ij}) - z(p\text{FA}_{ij}), \tag{S1}$$

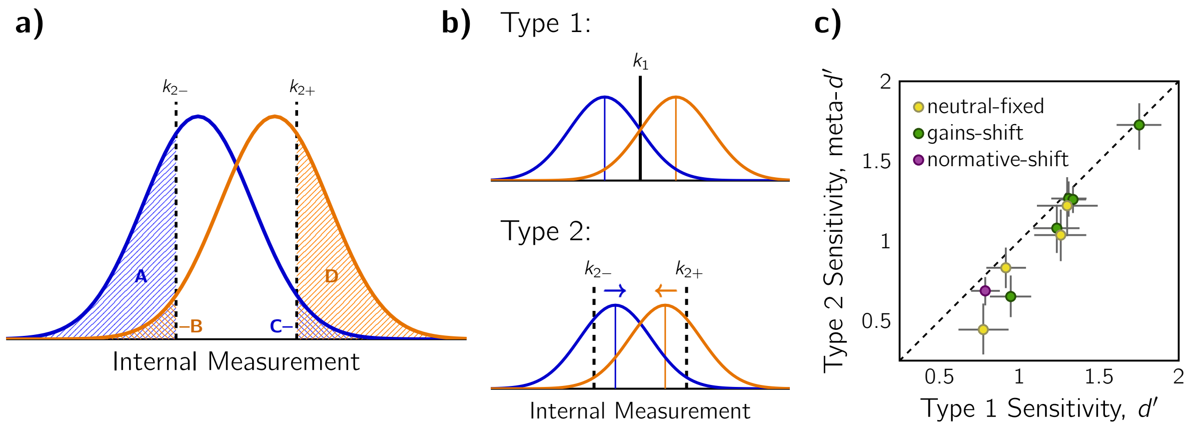


Figure S1: a) Depiction of example regions for the approximate meta-$d'$ calculation. Hatched regions correspond to the probability of a high-confidence judgment for the four possible pairings of stimulus and discrimination response. b) Example of greater sensitivity for perception (Type 1) than confidence (Type 2). In the standard SDT model, this corresponds to an inwards shift of the distributions for confidence. c) Contrast of $d'$ and meta-$d'$ results. Each data point is an observer, with 95% CIs derived from the posterior distribution of parameter estimates. Marker color indicates best-fitting Type 2 model. Dashed equality line is also shown for comparison.

where $p$H was the probability of selecting "right" when the stimulus was truly rightward tilted, $p$FA was the probability of selecting "right" when the stimulus was leftward tilted, and $z$ refers to the standard $z$-transform. In a similar fashion, we approximated the meta-$d'$ from the lower and upper confidence criteria, $k_{2-}$ and $k_{2+}$ respectively. These confidence criteria can be empirically calculated as per the standard method for deriving a criterion in Signal Detection Theory (SDT):

$$k_{2-} = \frac{1}{2}\big[z(p\mathrm{A}) + z(p\mathrm{B})\big] \tag{S2}$$

and

$$k_{2+} = \frac{1}{2}\big[z(p\mathrm{C}) + z(p\mathrm{D})\big]. \tag{S3}$$

The corresponding regions A-D are best demonstrated graphically (Figure S1a). To compute meta-$d'$, we used an average of two $d'$-like measurements, from the empirical upper and lower confidence bounds respectively:

$$\text{meta-}d'_{ij} = \frac{1}{2}\big[z(p\mathrm{A}_{ij}) - z(p\mathrm{B}_{ij}) + z(p\mathrm{D}_{ij}) - z(p\mathrm{C}_{ij})\big]. \tag{S4}$$

The concept behind computing a separate sensitivity parameter for confidence is that additional noise may have been applied to the internal measurement between the Type 1 and Type 2 decisions (Maniscalco and Lau, 2016). In the standard SDT framework, the variances of the distributions are fixed, and so the additional noise is modeled as a shift in distributions means (see Figure S1b). As such, we use the confidence bounds to estimate the relative separation of $p(x|S_L)$ and $p(x|S_R)$ with this additional metacognitive noise. These confidence bounds can then be represented in the original Type 1 space by a simple transformation

$$\frac{\text{meta-}d'}{d'} k_{2,\text{space2}} \rightarrow k_{2,\text{space1}}, \tag{S5}$$

as explained by Maniscalco and Lau (2012) and illustrated in Figure S1b.

In the hierarchical Bayesian model, each observation $j$ of $d'$ for participant $i$ was assumed to be drawn from a normally-distributed subject-specific prior,

$$d'_{ij} \sim \mathcal{N}(d'_i,\, \sigma_i^2), \tag{S6}$$

where $d'_i$ is the aggregate estimate of that participant's $d'$ for our next stage in modeling, and $\sigma_i^2$ is their sensitivity variance, capturing both noise in the calculation from a limited number of samples and sessional changes in sensitivity (e.g., attention, motivation). Similarly, we modeled the estimates of meta-$d'$ as

$$\text{meta-}d'_{ij} \sim \mathcal{N}(\text{meta-}d'_i,\, \sigma_i^2). \tag{S7}$$

Again, we have a subject-level estimate of sensitivity, meta-$d'_i$, for our modeling. The same variance parameter was used for both Type 1 and Type 2 estimates, because factors influencing noise in the observations are likely to be similar for both sensitivity measures. We also incorporated hyperpriors for both sensitivity measures, leveraging additional information we had about what to expect for these values. For $d'$, we used a normally-distributed hyperprior with a mean of 1.

$$d'_i \sim \mathcal{N}(1,\, \sigma^2_{\text{Type1}}), \tag{S8}$$

This decision was based on our expectations from the thresholding procedure, where the stimulus was adjusted to find $d' = 1$, and thus, on average, we expected this sensitivity for the observers in the main task. The population variance was $\sigma^2_{\text{Type1}}$. We also used the following hyperprior for meta-$d'$:

$$\text{meta-}d'_i \sim \mathcal{N}(0.8d'_i,\, \sigma^2_{\text{Type2}}). \tag{S9}$$

Based on previous results, we expected the meta-$d'$ of a participant to be, on average, about 80% of their $d'$ sensitivity measure (Maniscalco and Lau, 2012). Thus, the mean of the meta-$d'$ hyperprior was adjusted on a per-subject basis. There was a shared variance parameter, $\sigma^2_{\text{Type2}}$, representing variations in meta-cognition across participants in the same manner as $\sigma^2_{\text{Type1}}$. To ensure good model behavior, all free parameters had reasonable bounds imposed via a uniform prior either in addition to or in lieu of the other prior distributions described above: $[0, 3]$ for $d'_i$ and meta-$d'_i$, and $[0.1, 5]$ for $\sigma_i$, $\sigma_{\text{Type1}}$, and $\sigma_{\text{Type2}}$. The model was fit using custom-written scripts in the R and RStan programming languages (Carpenter et al., 2017), which implemented an MCMC fitting algorithm with 4000 iterations for each of 4 separate chains. The first half of the iterations were discarded as warmup. Parameter estimates and confidence intervals were calculated from the marginal posteriors (i.e., from the mean and percentile ranges of the samples).

The results of the model of Type 1 and Type 2 sensitivity are shown in Figure S1c. In general, there was greater sensitivity at the Type 1 level than at the Type 2 level, as expected (Maniscalco and Lau, 2012). The ratio of Type 2 to Type 1 sensitivity, also known as the *m-ratio* in the confidence literature (Fleming and Lau, 2014), was $0.86 \pm 0.04$ (mean±SEM). On average, participants' variability in $d'$ over sessions was $\hat{\sigma}_i = 0.19 \pm 0.02$ (mean±SEM). Across participants, we saw a variability in Type 1 sensitivity of $\hat{\sigma}_{\text{Type1}} = 0.37$ (95% CI: [0.23, 0.60] according to the posterior distribution of parameter fits), and at the Type 2 level, $\hat{\sigma}_{\text{Type2}} = 0.12$ (95% CI: [0.1, 0.35]).

#### 2 Multinomial Decision Model

Model fitting was performed in three sequential steps: (1) fitting of $d'$ and meta-$d'$, (2) Type 1 models, and (3) Type 2 models. In each case, the best-fitting parameters (and the best-fitting model in the Type 1 case) from one step were fixed while fitting models in the subsequent step. Fitting $d'$ and meta-$d'$ was explained in the previous section.

For Type 1 fits, we chose a dense grid of parameters, bias ($\gamma$) and between zero and three conservatism parameters ($\alpha$), with which to calculate the likelihood. The likelihood was a binomial across the two possible discrimination responses. We assumed a fixed lapse rate, $\lambda = 0.02$, for all participants, so

$$P(\text{data} \mid \theta) = \prod_{\text{stim} \in \{L,R\}} \prod_{\text{resp} \in \{\text{“}L\text{”},\text{“}R\text{”}\}} \Big(\lambda/2 \ + \ (1-\lambda)\, p(\text{resp} \mid \text{stim}, \theta)\Big)^{\mathcal{N}_{\text{resp,stim}}}, \qquad \text{(S10)}$$

where $\mathcal{N}_{\text{resp,stim}}$ is the number of trials in which that response was made for the discrimination of that stimulus.

The probability of a response is given by the corresponding area under the normal distribution, as in standard SDT. We fixed the variances of the internal response distributions to be 1, and positioned them based on the participant's sensitivity at locations $\pm d'/2$. Therefore, the probabilities for the correct responses, for example, were:

$$p(\text{“}L\text{”} \mid L) = \Phi\left(\gamma + k_1 + \frac{d'}{2}\right) \qquad \text{(S11)}$$

and

$$p(\text{“}R\text{”} \mid R) = 1 - \Phi\left(\gamma + k_1 - \frac{d'}{2}\right), \qquad \text{(S12)}$$

where $\Phi$ is the standard cumulative normal distribution. Note here that $k_1$ is calculated from $d'$ and $\alpha$ according to the Type 1 model.

The Type 2 fits inherited bias ($\gamma$) and various conservatism ($\alpha$) parameters from the Type 1 model fits. The $d'$ and meta-$d'$ values were inherited from the hierarchical $d'$ model fit. Thus, the counterfactual criterion $k_1^*$ was already fixed, and the Type 2 modeling involved only a single free parameter, $\delta$. Responses were modeled as a multinomial distribution with four possible responses to each stimulus, defined by the combination of the discrimination and confidence responses. We used the same lapse rate, but the probability of a particular random response was now halved because there were twice as many possible outcomes:

$$P(\text{data} \,|\, \delta) = \prod_{\text{stim}\in\{L,R\}} \prod_{\text{resp}\in\{\text{“}LH\text{”},\text{“}LL\text{”},\text{“}RH\text{”},\text{“}RL\text{”}\}} \left(\lambda/4 \ + \ (1-\lambda)\, p(\text{resp} \,|\, \text{stim}, \delta)\right)^{\mathcal{N}_{\text{resp,stim}}}. \tag{S13}$$

The probabilities of each response depend on the Type 2 criteria, for example:

$$p(\text{“}LH\text{”}|L) = \Phi\left(k_{2-} + \frac{d'}{2}\right) \tag{S14}$$

$$p(\text{“}LL\text{”}|L) = \Phi\left(k_1 + \frac{d'}{2}\right) - \Phi\left(k_{2-} + \frac{d'}{2}\right) \tag{S15}$$

$$p(\text{“}RL\text{”}|R) = \Phi\left(k_{2+} - \frac{d'}{2}\right) - \Phi\left(k_1 - \frac{d'}{2}\right) \tag{S16}$$

$$p(\text{“}RH\text{”}|R) = 1 - \Phi\left(k_{2+} - \frac{d'}{2}\right) \tag{S17}$$

$k_{2-}$ and $k_{2+}$ are the effective left and right confidence criteria respectively, and $\gamma$ was left out of these equations for readability. In the double-asymmetry conditions, it is possible for an observer's Type 1 criterion to be outside the intended symmetric bounds of the Type 2 criteria with a small enough $\delta$, as in Figure 1f. In this case, the effective $k_{2-}$ is actually equal to $k_1$. Concretely, this would happen if an observer was highly confident that the stimulus was right-tilted, but the potential rewards are so asymmetric that they respond left-tilted anyway. Because of the potential for these cases, $k_{2-}$ and $k_{2+}$ were not simply $k_1^* \pm \delta$, but rather

$$k_{2+} = \max(k_1, k_1^* + \delta) \tag{S18}$$

$$k_{2-} = \min(k_1, k_1^* - \delta). \tag{S19}$$

We used flat priors on all parameters, so we calculated model evidence by marginalizing across each dimension of the posterior.

$$p(\text{data}|M) = \int p(\text{data}|\theta, M)p(\theta)d\theta \tag{S20}$$

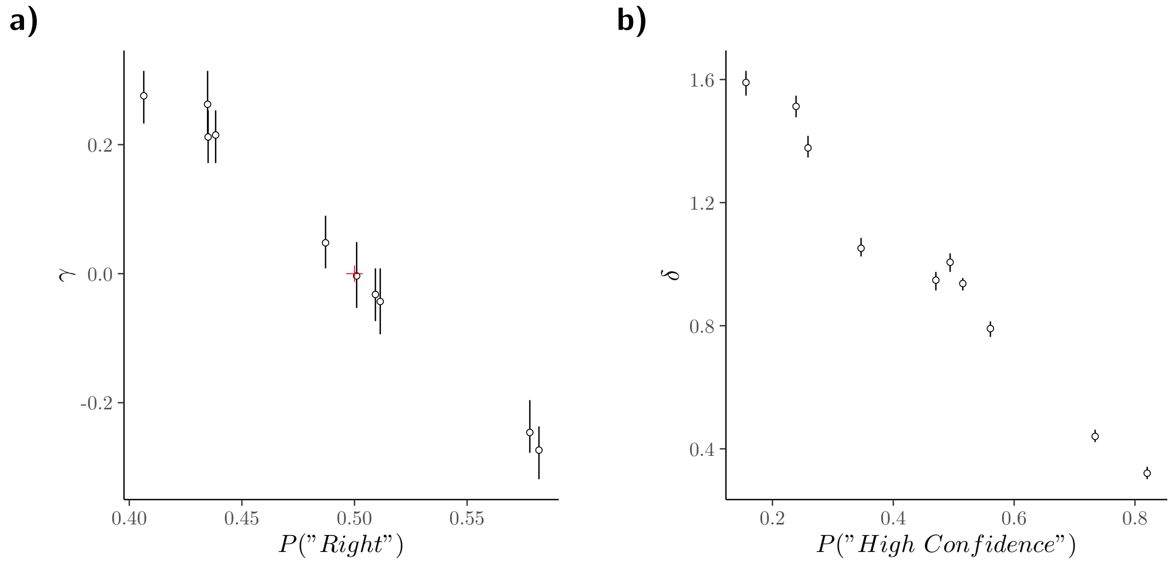


Figure S2: Checks on the fitted model parameters. a) Relationship between the bias in perceived vertical ($\gamma$) and the proportion of "right-tilt" judgments. Red cross: results for an unbiased observer. b) Relationship between the confidence criteria width parameter, $\delta$, and the proportion of "high confidence" judgments. Small $\delta$ leads to more high confidence reports (over-confidence). This predicted relationship is supported by the data. Error bars: 95% CIs from the posterior.

To do this, we numerically integrated the posterior of our parameter grid with a rectangular approximation by summing the volume of each grid element:

$$p(\text{data} \,|\, M) \approx \sum_{\theta} p(\text{data} \,|\, \theta, M)\, \Delta x_{\theta}, \tag{S21}$$

where $\Delta x_{\theta}$ is the product of step sizes for each dimension in the parameter grid. The model evidences for all models and all participants were used to compute the protected exceedance probability with the SPM12 Toolbox (Wellcome Trust Centre for Neuroimaging, London, UK) according to Rigoux et al. (2014).

### 3 Model Checks and Fits for All Subjects

Two of the model parameters make clear predictions about behavior. The fitted response bias parameter, $\gamma$, should be negatively correlated with the total proportion of trials the participants responded "right." Positive $\gamma$ values indicate a rightward tilted line is perceived as vertical, leading to fewer rightward responses overall. Figure S2a confirms this relationship ($r = -0.995, p < .0001$). The average bias is $\overline{\gamma} = .04 \pm .06$, with 70% of participants significantly biased according to the posterior parameter distribution. Also, $\delta$, half of the distance between the Type 2 criteria, should be inversely correlated with the proportion of "high confidence" reports; larger values of $\delta$ expand the low-confidence region (compare Figures 1e and f). This predicted relationship was obtained (Figure S2b; $r = -0.986, p < .0001$; $\overline{\delta} = 1.00 \pm 0.13$). These predictions are not trivial: idiosyncratic biases in one condition may disappear or reverse on a subsequent day in the inverse condition. Nevertheless, we find that the $\gamma$ and $\delta$ parameters are meaningfully capturing patterns of behavior.

The following figures show the results of all subjects in the style of Figures 4 and 5 of the main paper.

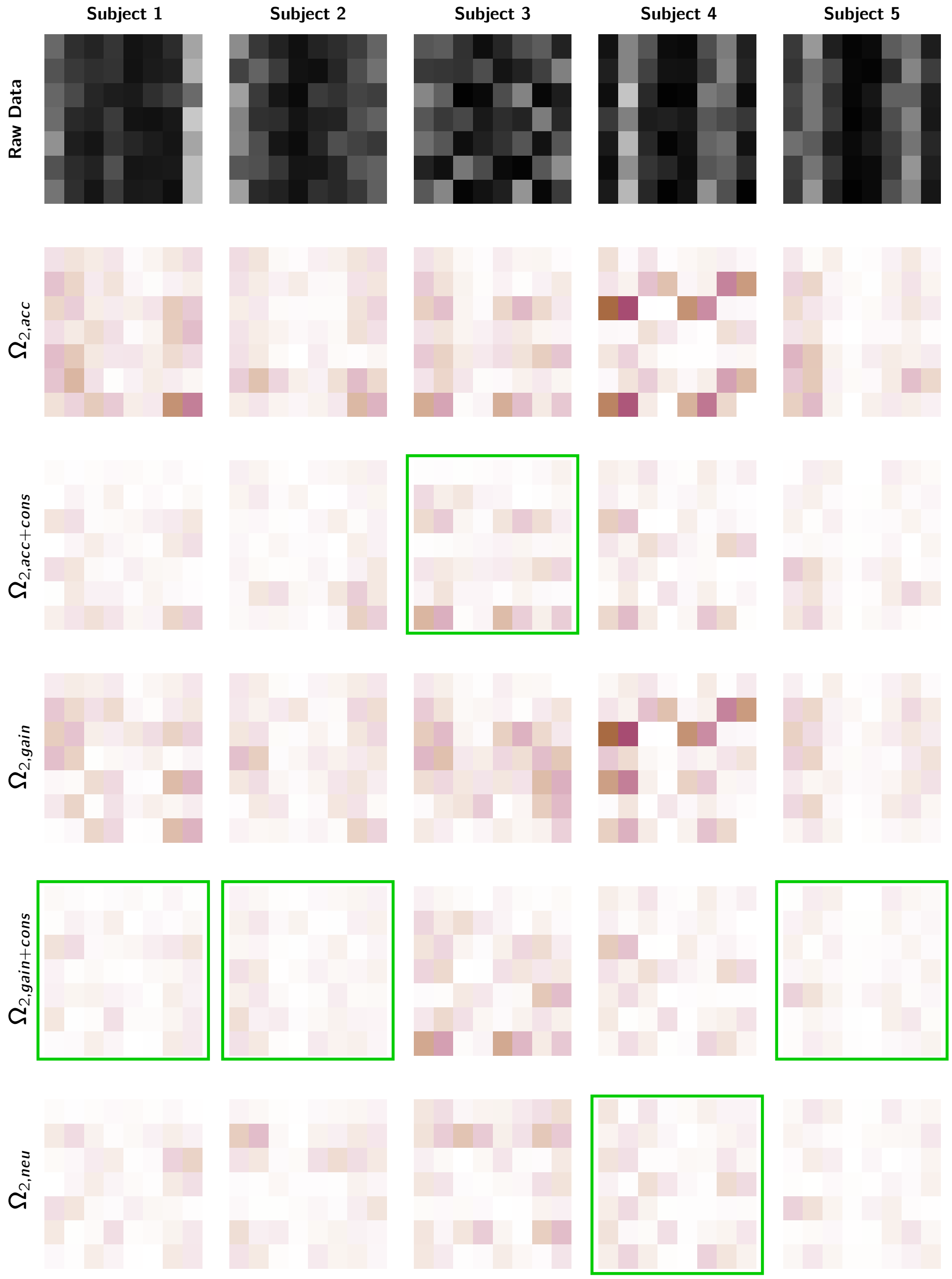


Figure S3: Raw and predicted response rates for participants 1-5. Grids are formed from the seven conditions (rows) and the eight possible stimulus-response-confidence combinations (columns). Condition order: (1) full symmetry, (2) single asymmetry ($p(R) = .75$), (3) single asymmetry ($p(R) = .25$), (4) single asymmetry ($V_R : V_L = 4 : 2$), (5) single asymmetry ($V_R : V_L = 2 : 4$), (6) double asymmetry ($p(R) = .75, V_R : V_L = 2 : 4$), (7) double asymmetry ($p(R) = .25, V_R : V_L = 4 : 2$). Fill: proportion of trials for that condition and stimulus that have that combination of response and confidence. Top row: Raw response rates. Subsequent rows: difference between raw and predicted response rates as per each model. Green boxes: winning models.

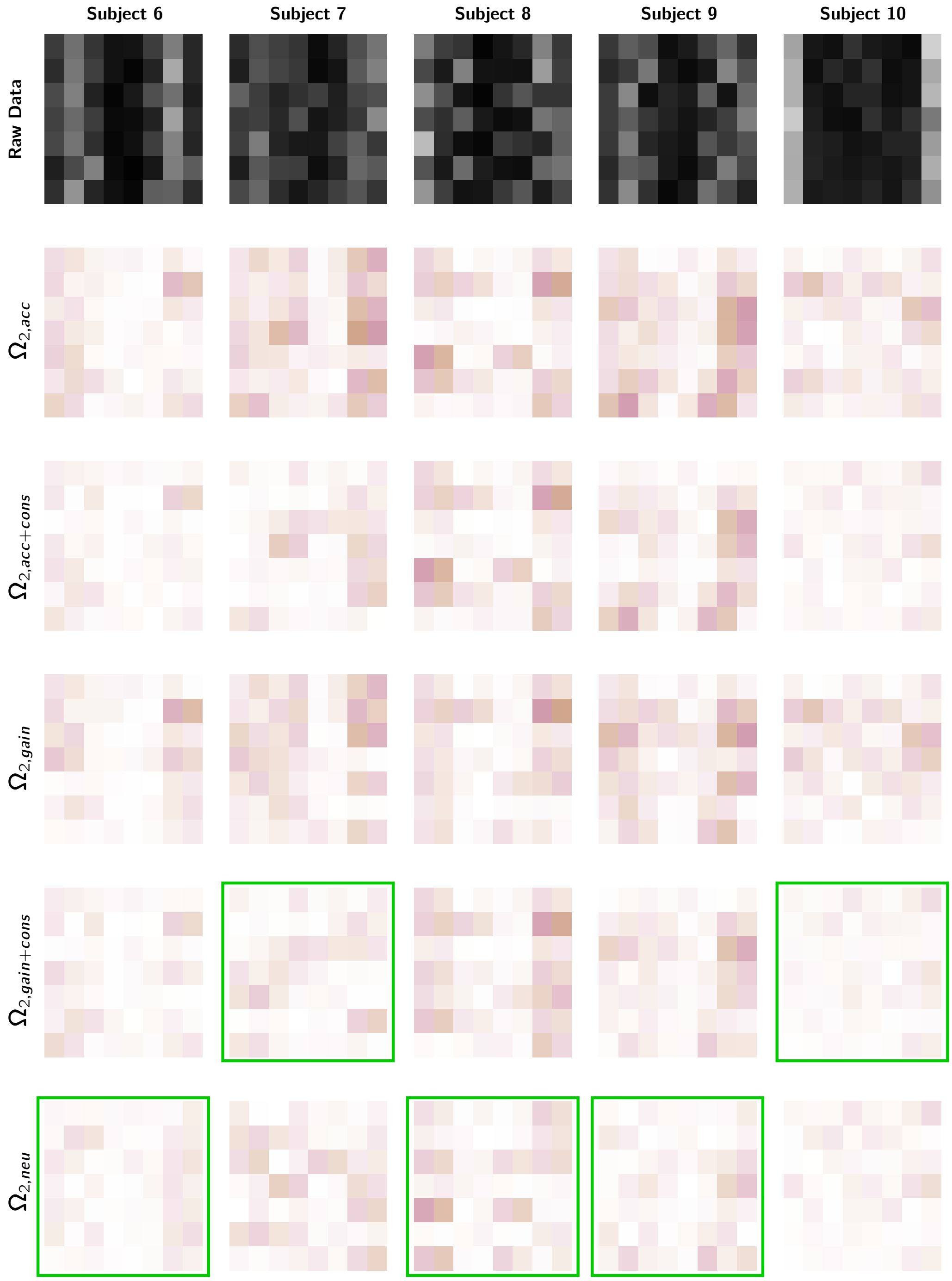


Figure S4: Raw proportions of subjects 6-10 in the style of Figure S3.

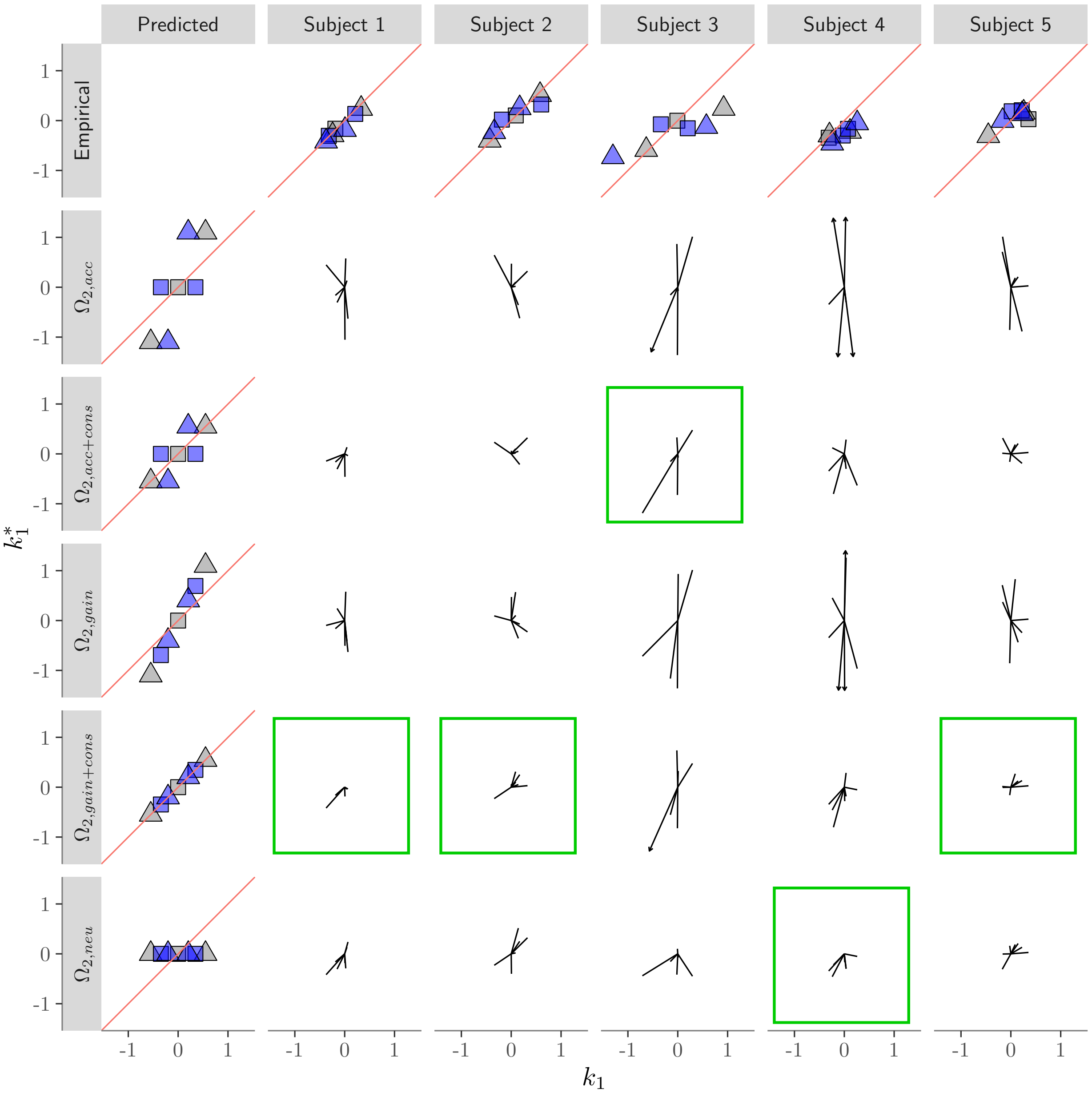


Figure S5: Comparison of the empirical and predicted $k_1$ and $k_1^*$ for participants 1-5. Top row: empirical criteria. $k_1^*$ was calculated as the midpoint between the two empirical $k_2$ (see Figure S1 for $k_2$ calculation details). Left column: predicted relationship between the Type 1 and Type 2 criteria ($d' = 1$; all $\Omega_{1,1\alpha}$ with $\alpha = 0.5$). Grey and square symbols: symmetry conditions. Triangles: prior asymmetry. Blue symbols: payoff asymmetry. Polar plots: residuals between empirical data and model prediction based on best-fitting parameters, plotted as vectors. Arrowheads: residuals greater than plot bounds.

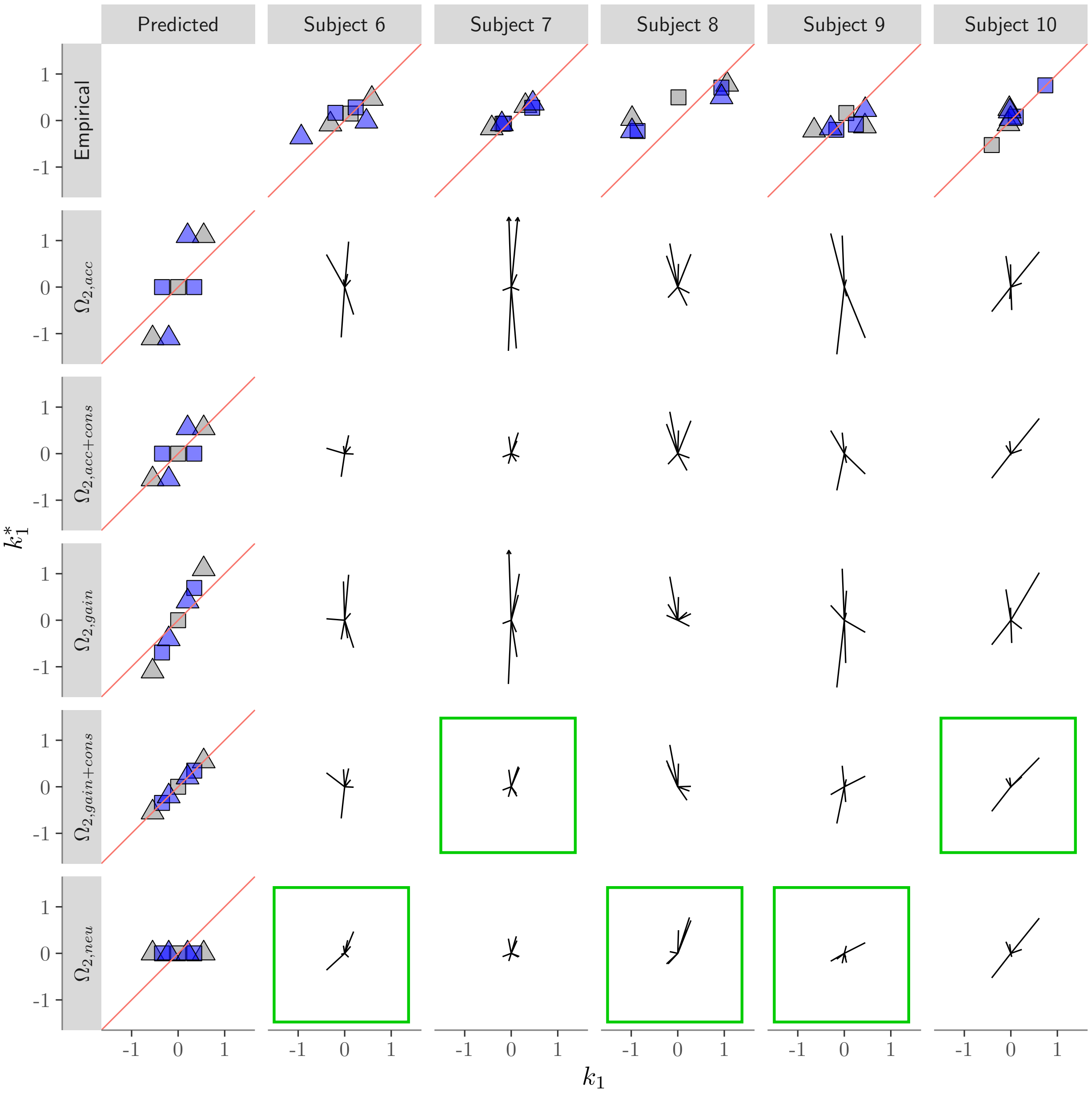


Figure S6: Comparison of $k_1$ and $k_1^*$ for participants 6-10 in the style of Figure S5.

### 4 Gains-Accuracy Trade-off Strategy and Conservatism

Here, we show how the gain-accuracy trade-off strategy of Maddox and Bohil (1998) is equivalent to the $\Omega_{1,2\alpha}$ model. The gain-accuracy trade-off strategy can be expressed mathematically as a weighted sum between the gain-maximizing criterion, $k_{opt}$, and the accuracy-maximizing criterion, $k_p$, with weight $w$ ($0 \leq w \leq 1$). We also applied a single general conservatism parameter in this weighting strategy, which can be thought of as acting on each separate component or equivalently to the sum of the components. A simple rearrangement shows how these two models are equivalent:

$$\begin{aligned} \alpha_v k_v + \alpha_p k_p &= w\alpha k_{opt} + (1-w)\alpha k_p \\ &= \alpha(wk_v + wk_p + k_p - wk_p) \\ &= \alpha(wk_v + k_p) \\ &= \alpha w k_v + \alpha k_p \end{aligned} \tag{S22}$$

Therefore, we find that different degrees of conservatism for priors than payoffs can arise as a result of weight values less than 1. Specifically, the weight value contributes to an increase in a general level of conservatism, $\alpha_v = \alpha w$ and $\alpha_p = \alpha$, where the constraint $w \leq 1$ ensures that $\alpha_v \leq \alpha_p$. If $w = 1$, then $\alpha_v = \alpha_p = \alpha$, which is the single conservatism model $\Omega_{1,1\alpha}$.
